# Spray-induced gene silencing for disease control is dependent on the efficiency of pathogen RNA uptake

**DOI:** 10.1101/2021.02.01.429265

**Authors:** Lulu Qiao, Chi Lan, Luca Capriotti, Audrey Ah-Fong, Jonatan Nino Sanchez, Rachael Hamby, Jens Heller, Hongwei Zhao, N. Louise Glass, Howard S. Judelson, Bruno Mezzetti, Dongdong Niu, Hailing Jin

**Affiliations:** College of Plant Protection, Nanjing Agricultural University, Nanjing 210095, China; Key Laboratory of Integrated Management of Crop Diseases and Pests (Ministry of Education), Nanjing 210095, China; Department of Microbiology & Plant Pathology, Center for Plant Cell Biology, Institute for Integrative Genome Biology, University of California, Riverside, California 92521, USA; Department of Agricultural, Food and Environmental Sciences, Marche Polytechnic University, 60131 Ancona, Italy; Department of Plant and Microbial Biology, University of California, Berkeley, California 94720, USA; Environmental Genomics and Systems Biology Division, The Lawrence Berkeley National Laboratory, Berkeley, California 94720, USA

**Keywords:** spray-induced gene silencing (SIGS), small RNA, RNA interference, double-stranded RNA (dsRNA), uptake efficiency

## Abstract

Recent discoveries show that fungi can take up environmental RNA, which can then silence fungal genes through environmental RNA interference. This discovery prompted the development of <u>S</u>pray-<u>I</u>nduced <u>G</u>ene <u>S</u>ilencing (SIGS) for plant disease management. In this study, we aimed to determine the efficacy of SIGS across a variety of eukaryotic microbes. We first examined the efficiency of RNA uptake in multiple pathogenic and non-pathogenic fungi, and an oomycete pathogen. We observed efficient double-stranded RNA (dsRNA) uptake in the fungal plant pathogens *Botrytis cinerea*, *Sclerotinia sclerotiorum*, *Rhizoctonia solani*, *Aspergillus niger*, and *Verticillium dahliae*, but no uptake in *Colletotrichum gloeosporioides*, and weak uptake in a beneficial fungus, *Trichoderma virens*. For the oomycete plant pathogen, *Phytophthora infestans*, RNA uptake was limited, and varied across different cell types and developmental stages. Topical application of dsRNA targeting virulence-related genes in the pathogens with high RNA uptake efficiency significantly inhibited plant disease symptoms, whereas the application of dsRNA in pathogens with low RNA uptake efficiency did not suppress infection. Our results have revealed that dsRNA uptake efficiencies vary across eukaryotic microbe species and cell types. The success of SIGS for plant disease management can largely be determined by the pathogen RNA uptake efficiency.

## Introduction

Crop protection at both the pre- and post-harvest levels is critical for long-term food security. Fungal and oomycete diseases pose a serious threat to crop production worldwide (Bebber and Gurr, 2015). At present, agricultural crops rely almost exclusively on fungicides to control disease, resulting in pesticide residues that often threaten human health and the environment. Further, resistant strains of fungi have been identified against every major fungicide used in agricultural applications (Fisher et al., 2018). Thus, there is an urgent need to develop eco-friendly, effective solutions to control plant diseases. The development of disease control strategies using biomolecules (nucleic acids, lipids, sugars and proteins) that exploit naturally occurring pathways may help circumvent the use of chemical pesticides.

Recent studies have discovered a novel mechanism of communication between plants and their pathogens, termed cross-kingdom RNA interference (RNAi), a phenomenon in which fungi send small RNAs (sRNAs) into host plants to silence host immune response genes and plants also send sRNAs packaged in extracellular vesicles into their fungal pathogens to silence virulence-related genes (Cai et al., 2018b; Huang et al., 2019; Weiberg et al., 2013; Zeng et al., 2019). Cross-kingdom RNAi can be utilized in plant protection strategies by genetically modifying host plants to express sRNAs or double-stranded RNAs (dsRNAs) that target pathogen virulence-related genes, a technology called host-induced gene silencing (HIGS) (Cai et al., 2019; Song and Thomma, 2018; Wang et al., 2017a; Zhang et al., 2016). These transgene-derived artificial RNAs can be delivered from plants to pathogens and pests, including fungi (Cai et al., 2018a; Nowara et al., 2010; Qi et al., 2019; Xu et al., 2018), nematodes (Chen et al., 2015; Shivakumara et al., 2017), oomycetes (Govindarajulu et al., 2015; Jahan et al., 2015), and insects (Abdellatef et al., 2015; Baum et al., 2007; Liu et al., 2019), inducing gene silencing in *trans* and conferring disease resistance in the host plants. However, the application of HIGS, is limited due to the lack of efficient transformation techniques in many crop plants, including many fruit trees, vegetables, and flowers (Capriotti et al., 2020; Koch and Kogel, 2014). Furthermore, many consumers still resist genetically modified (GMO) products, which are currently banned in European agricultural production (Kleter et al., 2018; Tsatsakis et al., 2017; Wunderlich and Gatto, 2015). Therefore, innovative non-GMO plant protection strategies are highly desired.

Previous work has found that fungal pathogens *Botrytis cinerea* and *Fusarium graminearum* can efficiently take up environmental dsRNAs, which are then processed into sRNAs and induce the silencing of pathogen genes with complementary sequences (Koch et al., 2016; Wang et al., 2016). These discoveries prompted the development of an innovative crop protection strategy, <u>S</u>pray-<u>I</u>nduced <u>G</u>ene <u>S</u>ilencing (SIGS). As a non-GMO alternative to HIGS, SIGS inhibits pathogen infection by topical application of dsRNA or sRNA molecules onto plants to silence pathogen virulence-related genes (Song et al., 2018; Wang and Jin, 2017). SIGS has been successfully utilized to prevent some fungal diseases. Topical application of dsRNAs and sRNAs that target the key components of the RNAi machinery, *Dicer-like* (*DCL*)*1*, and *DCL2* genes of the fungal pathogen *Botrytis cinerea*, inhibits grey mold disease on fruits, vegetables, and flowers (Wang et al., 2016). When applied to barley leaves, dsRNA designed to target fungal cytochrome P450 inhibited the growth of the fungal pathogen *Fusarium graminearum* (Koch et al., 2016). Additionally, dsRNA sprays can inhibit *Botrytis cinerea* and *Sclerotinia sclerotiorum* growing on *Brassica napus* (McLoughlin et al., 2018). However, it is still not clear what determines SIGS efficacy and how effective SIGS is against a wide variety of eukaryotic pathogens. Establishing the effectiveness of SIGS across a wide range of pathogens is a critical next step in the development of this technology.

For SIGS technology to be effective, it is crucial that fungi can take up RNA from the environment. Here, we investigated the environmental RNA uptake efficiency of various plant pathogenic fungi. These included *S. sclerotiorum*, a necrotrophic plant pathogen which infects over 400 plant species worldwide, principally vegetables and ornamental plants (Bolton et al., 2006; Kabbage et al., 2015); *Aspergillus niger*, a haploid filamentous fungus that can contaminate fruits, cereal grains, before or after harvest, and some animal products directly by fungal growth or indirectly through the production of mycotoxins (Hocking et al., 2007; Wilson et al., 2002); *Rhizoctonia solani*, a well-known soil-borne fungus that causes a wide range of significant crop diseases (Zhou et al., 2016a); *Verticillium dahliae*, a soil-borne fungus that causes wilt disease in a wide range of plant species (Klosterman et al., 2009; Luo et al., 2014); and *Colletotrichum gloeosporioides*, which causes anthracnose, a disease characterized by sunken necrotic lesions, which can infect fruits, flowers, leaves, petioles, stolons, and crowns (Zhou et al., 2016b). In addition to various pathogenic fungi, we also investigated the RNA uptake efficiency of a non-pathogenic fungus, *Trichoderma virens*, which is a biocontrol agent of plant diseases (Contreras-Cornejo et al., 2011; Yang et al., 2016). Finally, we investigated the RNA uptake ability of a non-fungal eukaryotic pathogen, the oomycete *Phytophthora infestans*, which is the causal agent of late blight disease that threatens tomato and potato crops worldwide (Nowicki et al., 2012).

In this study, we found that many eukaryotic microbes can take up RNA from the environment but with different efficiencies. *B. cinerea* and *S. sclerotiorum*, as shown previously (McLoughlin et al., 2018; Wang et al., 2016), *R. solani*, *A. niger*, and *V. dahliae* could take up environmental RNA with high efficiency, whereas RNA uptake was modest in *T. virens*, undetectable in *C. gloeosporioides*, and limited in *P. infestans.* Furthermore, we found that topical application of dsRNAs targeting virulence-related pathogen genes could suppress disease caused by the pathogens that have high RNA uptake efficiency. In comparison, SIGS did not inhibit infection of *C. gloeosporioides* due to the lack of environmental RNA uptake. Finally, to examine the longevity of dsRNA-conferred protection on pre-harvesting crops, we challenged tomato leaves with *B. cinerea* over a time-course post dsRNA treatment. In summary, we found that most fungi tested can take up dsRNA, and efficient dsRNA uptake in pathogens is essential for success of SIGS in crop protection.

## Results

### RNA uptake efficiency varies across fungal species

To determine whether dsRNA uptake by fungi is a common phenomenon, we first tested the efficiency of dsRNA uptake across different fungal species, including pathogens from different classes of Ascomycota: *B. cinerea* and *S. sclerotiorum* from the Leotiomycete class, *A. niger* from the Eurotiomycete class, *V. dahliae* from the Sordariomycetes class, and a non-pathogenic fungus, *T. virens* from the Sordariomycetes class of Ascomycota (Fig. S1), a common soil microbe that is usually part of healthy root ecosystems that benefit plant growth (Contreras-Cornejo et al., 2009). We also tested a rice pathogen *R. solani* from the Basidiomycota division. To determine RNA uptake efficiency, we treated fungal cells with fluorescein-labeled *YFP*-dsRNA. These fungal cells had no obvious autofluorescence (Fig. S2), and therefore, there will be no interference with the fluorescence signal from the fluorescein-labeled dsRNA. Micrococcal nuclease (MNase) treatment was performed 30 minutes before microscopy analysis to remove any labeled dsRNAs that were outside the fungal cells. We found that different fungal species had different rates and efficiencies of dsRNA uptake (Fig. 1 and Fig. S3). Fluorescence signals were observed as early as 6 hours post dsRNA treatment (hpt) in *B. cinerea, S. sclerotiorum* and *R. solani* (Fig. S3), all of which showed very efficient RNA uptake (Fig. 1a). In *A. niger,* an intense fluorescence signal was observable 10 hpt, and in *V. dahliae* 12 hpt (Fig. 1a and Fig. S3). However, no fluorescence signal was detectable in *C. gloeosporioides* even 24 hpt, suggesting that *C. gloesporioides* cannot take up environmental dsRNA (Fig. 1b-c and Fig. S3). Further, *T. virens* mycelium showed only very faint RNA fluorescence signals even at 30 hpt (Fig. 1b and Fig. S3).

**Figure 1.**
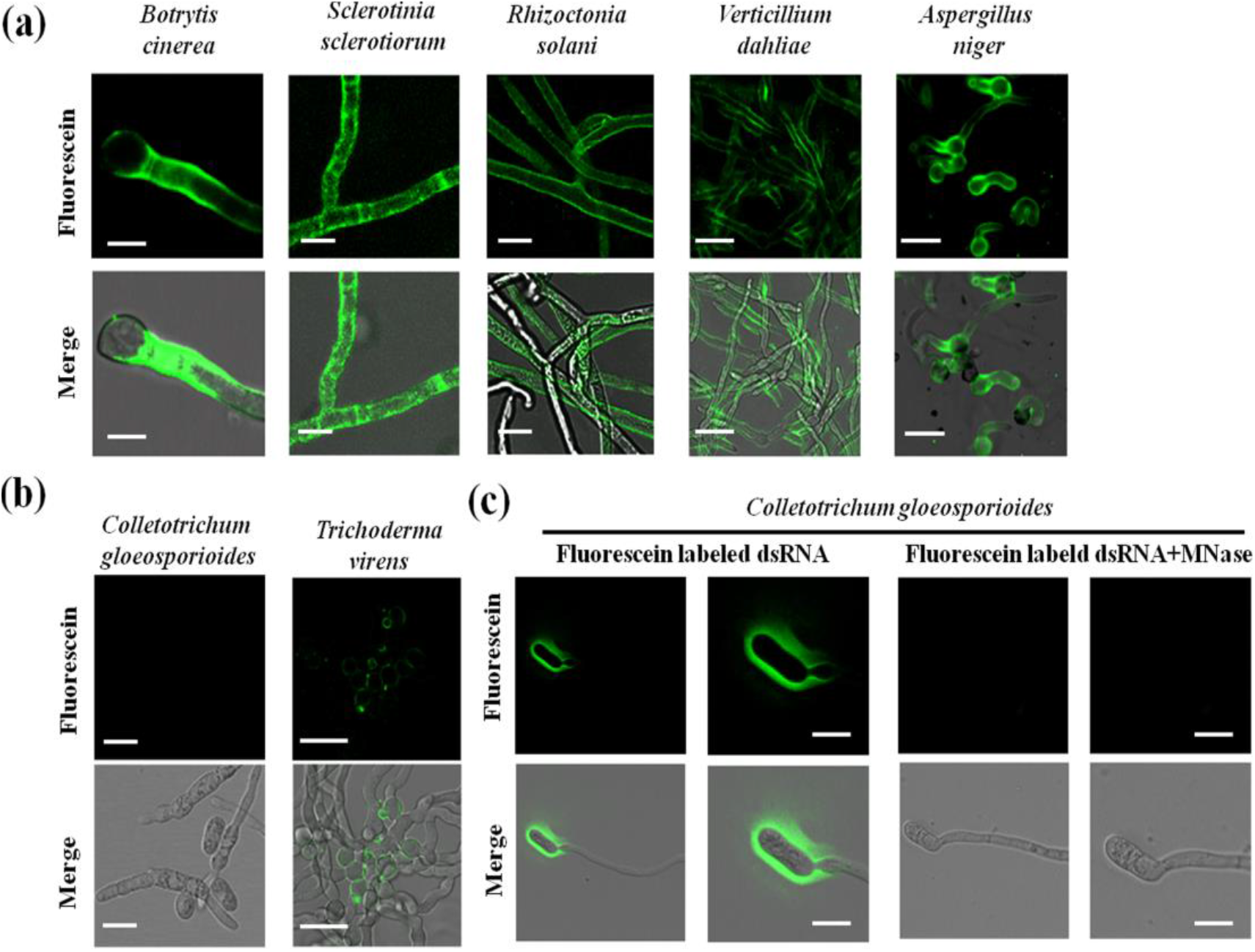
Examination of dsRNA uptake efficiencies in multiple fungi using fluorescein-labeled *YFP*-dsRNA. **(a) (b)** Fluorescein-labeled *YFP*-dsRNA was added to the spores of *B. cinerea*, *V. dahliae*, *A. niger*, *C. gloeosporoides and T. virens,* and to the hyphae of *S. sclerotiorum* and *R. solani*. Micrococcal nuclease (MNase) treatment was performed 30 min before acquiring images using the confocal microscopy laser scanner (CMLS). Fluorescence signals were detected inside of *B. cinerea, S. sclerotiorum*, *R. solani, A. niger,* and *V. dahliae* cells, but not *C. gloeosporoides* cells. A weak signal was observed in *T. virens* cells. Pictures were taken at 10 hours post treatment (hpt) for *B. cinerea, S. sclerotiorum*, and *R. solani, A. niger,* and *V. dahliae,* and at 24 hpt for *C. gloeosporoides* and *T. virens*. Scale bars = 20 μm except for *B. cinerea* image (Scale bars = 10 μm). **(c)** *C. gloeosporioides* spores are unable to take up dsRNA. Fluorescence signals were only visible on the outer surface of *C. gloeosporioides* cells before MNase treatment, but disappeared after MNase treatment. Scale bars = 15 μm.

To further confirm that the dsRNA entered the cytoplasm of the fungal cells, the fluorescein-labeled dsRNA was added to the liquid culture of *S. sclerotiorum* mycelium, which was subjected to protoplast preparation after culturing for 48 hours. Fluorescence signals were clearly observed within *S. sclerotiorum* protoplasts after MNase treatment (Fig. S4). Overall, we found that *B. cinerea, S. sclerotiorum*, *R. solani*, *V. dahliae*, and *A. niger* took up external dsRNA efficiently although *V. dahliae*, and *A. niger* were at a slower rate (Fig. 1 and Figs. S3, S5), whereas *C. gloeosporioides* showed no detectable dsRNA uptake and the non-pathogenic fungus *T. virens* displayed weak and slow dsRNA uptake (Fig. 1 and Figs. S3, S5). Thus, dsRNA uptake exhibited different efficiencies across diverse species of fungi.

### Topical application of dsRNA targeting vesicle-trafficking pathway genes inhibited the virulence of *B. cinerea, S. sclerotiorum* and *A. niger*

Our recent work showed that *Botrytis* single deletion mutant strains of vesicle-trafficking pathway genes v*acuolar protein sorting 51* (*VPS51*), *dynactin* (*DCTN1*), and *suppressor of actin* (*SAC1*) had greatly reduced virulence on *Arabidopsis thaliana* leaves (Cai et al., 2018b). To validate whether SIGS of vesicle-trafficking pathway genes could control grey mold disease, we first generated a *Bc-VPS51+DCTN1+SAC1*-dsRNA construct by integrating non-conserved fragments of *Bc-VPS51* (169 bp), *Bc-DCTN1* (179 bp), and *BcSAC1* (168 bp) via overlap PCR and transcribing the construct *in vitro* from both the 5’ and 3’ ends under T7 promoter (Table S2). In all dsRNA constructs throughout this paper, non-conserved regions are selected to avoid off-target effects in host plants or beneficial microbes. *Bc-VPS51+DCTN1+SAC1* or *Bc-DCL1/2* dsRNA (20 μl of 20 ng μl^−1^ in RNase-free water) were separately applied to lettuce leaves, tomato fruits, rose petals, and grape berries. After dsRNA treatment, plant materials were then inoculated with *B. cinerea* at the dsRNA treated area. All plant materials treated with *Bc-VPS51+DCTN1+SAC1-*dsRNA showed a reduction in disease symptoms as compared to the controls treated with water or *YFP*-dsRNA (Fig. 2a, b). The reduction of the infected areas was comparable to infection phenotypes after *Bc-DCL1/2*-dsRNA treatment (Fig. 2a, b), which was used as a positive control as previous work demonstrated Bc-DCL1/2-dsRNA was effective in reducing Botrytis virulence (Wang et al., 2016). Moreover, the mRNA expression levels of *BcVPS51*, *BcDCTN1*, and *BcSAC1* mRNA were reduced in *B. cinerea* treated with *Bc-VPS51+DCTN1+SAC1*-dsRNA (Fig. 2c).

**Figure 2.**
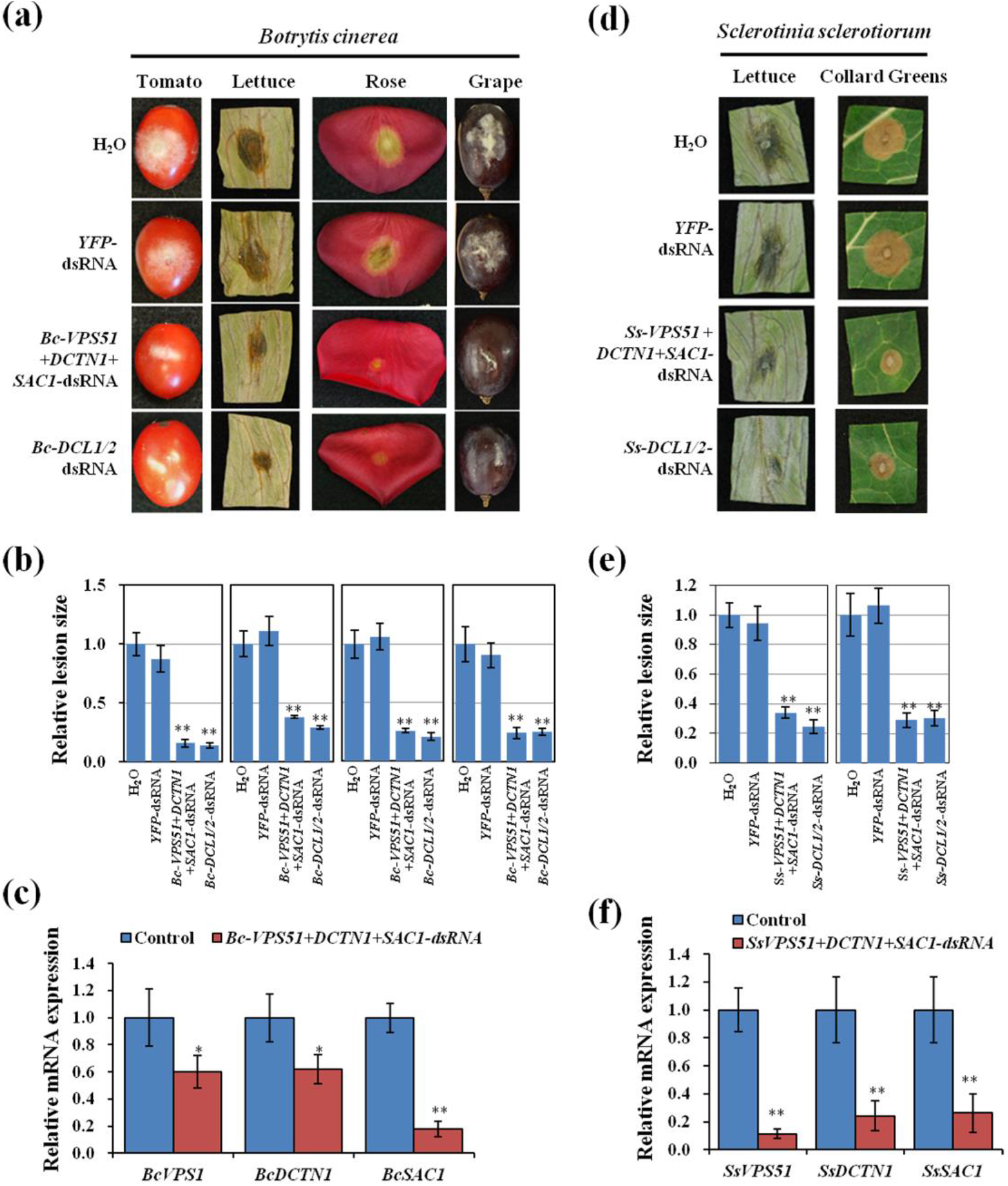
Topical application of pathogen gene-targeting dsRNAs inhibited the virulence of *B. cinerea* and *S. sclerotiorum*. **(a)** Tomato fruits, lettuce leaves, rose petals, and grape berries were inoculated with *B. cinerea* spores after treating with controls (water or *YFP*-dsRNA) or *Bc-VPS51+DCTN1+SAC1-*dsRNA or *Bc-DCL1/2*-dsRNA (20 ng μl^−1^). **(b)** The relative lesion sizes were measured 3 days post-inoculation (dpi) on lettuce leaves, rose petals, and 5 dpi on grapes and tomato fruits using ImageJ software, respectively. Error bars indicate the SD of 10 samples and three biological repeats were conducted for the relative lesion sizes. Statistical significance (Student’s t-test): **, *P* < 0.01. **(c)** qRT-PCR analysis of *Bc-VPS51*, *Bc-DCTN1*, and *Bc-SAC1* expression in *Bc-VPS51+DCTN1+SAC1-*dsRNA treated *B. cinerea* cells. Error bars represent the SD from three technical replicates. Statistical significance (Student’s t-test): *, *P* < 0.05; **, *P* < 0.01 between the treatment and the water control. **(d)** Lettuce and collard green leaves were inoculated with *S. sclerotiorum* mycelium plugs after treating with controls (water or *YFP*-dsRNA) or *Ss-VPS51+DCTN1+SAC1-* and *Ss-DCL1/2*-dsRNA (40 ng μl^−1^). Pictures were taken at 3 dpi. **(e)** The relative lesion sizes were measured at 3 dpi for lettuce and collard greens using ImageJ software. Error bars indicate the standard deviations (SD) of 10 samples. Asterisks (**) indicate statistically significant differences (*P* < 0.01, Student’s t-test) between the treatment and the water control. **(f)** The mRNA expression levels of *SsVPS51*, *SsDCTN1*, and *SsSAC1* were detected in *Ss-VPS51+DCTN1+SAC1*-dsRNA-treated *S*. *sclerotiorum* by qRT-PCR analysis. Asterisks (**) indicate statistically significant differences (*P* < 0.01, Student’s t-test) between the treatment and the water control. Similar results were observed from three biological replicates in (c) and (f).

*S. sclerotiorum* is closely related to *B. cinerea.* To test whether *VPS51, DCTN1* and *SAC1* may also play an essential role for *S. sclerotiorum* pathogenicity, we knocked out *SsVPS51*, *SsDCTN1*, and *SsSAC1* genes by homologous recombination. *Ss*-*Δvps51*, *Ss*-*Δdctn1*, and *Ss*-*Δsac1* mutant strains showed a reduction in development and mycelium growth after 3 days of culture and reduced sclerotia size on PDA media in comparison to the wild-type (WT) strain (Fig. S6a, b). Most importantly, the *Ss*-*Δvps51*, *Ss*-*Δdctn1*, and *Ss*-*Δsac1* knockout mutants exhibited reduced virulence on lettuce and collard green leaves in comparison to WT (Fig. S6c, d). Thus, *S. sclerotiorum VPS51*, *DCTN1*, and *SAC1* are important for the pathogenicity of *S. sclerotiorum* and should be appropriate targets for SIGS.

Next, we generated *S. sclerotiorum VPS51+DCTN1*+*SAC1*-dsRNA by integrating *SsVPS51* (230 bp), *SsDCTN1* (196 bp), and *SsSAC1* (194 bp) via overlap PCR and performed *in vitro* transcription from both the 5’ and 3’ ends (Table S2). The *Ss-VPS51+DCTN1+SAC1-*dsRNA (20 μl of 40 ng μl^−1^) was externally applied to lettuce and collard green leaves. After dsRNA application, plant material was challenged with *S. sclerotiorum*. All plants treated with *Ss-VPS51+DCTN1+SAC1*-dsRNA showed reduced disease symptoms and lesion size compared to water and *YFP*-dsRNA treated controls (Fig. 2d, e). Moreover, the mRNA expression levels of *SsVPS51*, *SsDCTN1*, and *SsSAC1* were also reduced in *S. sclerotiorum* (Fig. 2f). Furthermore, *Ss-DCL1+DCL2-*dsRNA application could also inhibit *S. sclerotiorum* infection, suggesting that DCL proteins are important for the virulence of *S. sclerotiorum*. Thus, externally applied *Ss-VPS51+DCTN1+SAC1*-dsRNA or *Ss-DCL1+DCL2-*dsRNA protected plants from *S. sclerotiorum* invasion.

*Aspergillus niger* can cause postharvest diseases in many fruits and vegetables (Liu et al., 2017). The *A. niger* genome encodes the vesicle trafficking pathway genes *VPS51*, *DCTN1* and *SAC1. A. niger VPS51+DCTN1+SAC1*-dsRNA was generated accordingly. Furthermore, previous research suggested that a probable exo-polygalacturonase b (*pgxB*) is an *A. niger* virulence gene, which plays an important role in the infection process on the fruit (Liu et al., 2017). dsRNA targeting a non-conserved region of *pgxB* (219 bp) was also generated. Topical application of the *VPS51+DCTN1+SAC1*-dsRNA or the *pgxB-* dsRNA onto the surface of tomato, apple and grape fruits led to much weaker fungal growth and disease symptoms, resulting in significantly smaller lesions than the control samples treated with *YFP*-dsRNA or water (Fig. 3a, b). In addition, the mRNA expression levels of the target genes were reduced in *A. niger* after *pgxB*- and *VPS51+DCTN1+SAC1*-dsRNA treatment (Fig. 3c). Thus, external application of *VPS51+DCTN1+SAC1-*dsRNA inhibits infection of *Botrytis cinerea, S. sclerotiorum* and *A. niger.*

**Figure 3.**
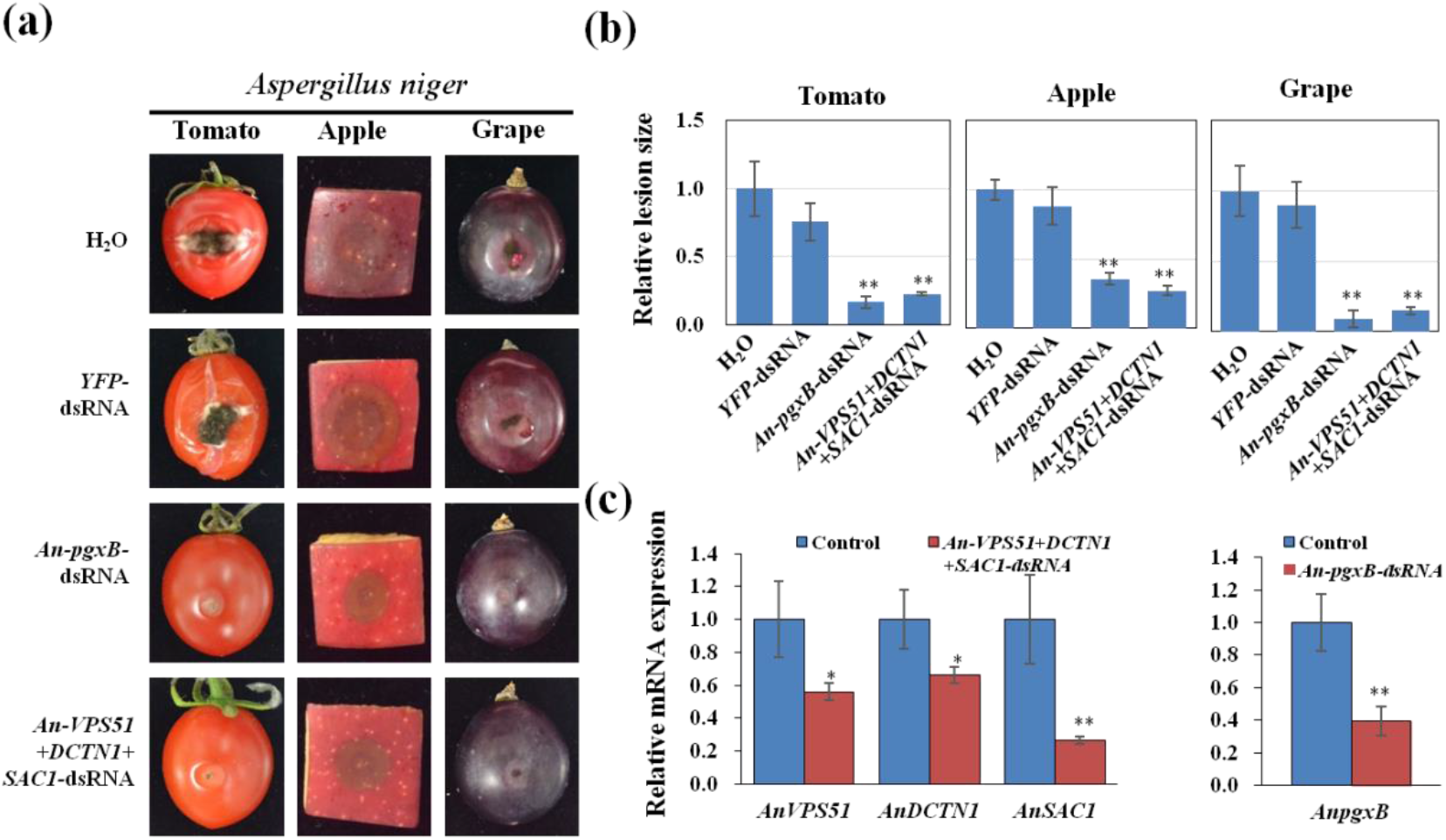
Topical application of pathogen gene-targeting dsRNAs inhibited the *A. niger* virulence. **(a)** Tomatoes, apples, and grapes were inoculated with *A. niger* spores after treating with controls (water or *YFP*-dsRNA), *An-pgxB-* or *An-VPS51+ DCTN1+SAC1*-dsRNA (20 ng μl^−1^). Pictures were taken at 5 dpi. **(b)** The relative lesion sizes were measured at 5 dpi using ImageJ software. Error bars indicate the SD of 10 samples. **(c)** The mRNA expression levels of *AnVPS51*, *AnDCTN1*, *AnSAC1* and *AnpgxB* were detected in *An-VPS51+DCTN1+SAC1*-dsRNA- or *An-pgxB*-treated *A. niger,* respectively. Error bars indicate the SD of three technical replicates. Statistical significance (Student’s t-test) between the treatment and the water control: *, *P* < 0.05; **, *P* < 0.01. Similar results were observed from three biological replicates.

### Topical application of dsRNA inhibited the virulence of *R. solani*

Rice sheath blight caused by *R. solani* is one of the most devastating fungal diseases of rice (Oryza sativa) worldwide. We identified the vesicle trafficking pathway genes *DCTN1* and *SAC1* in the *R. solani* genome (Zheng et al., 2013), but *VPS51* was not identified. To analyze whether SIGS could be adapted to control sheath blight, we generated *Rs-DCTN1+SAC1*-dsRNA containing two segments from non-conserved regions of *Rs-DCTN1*, and *Rs-SAC1* and applied it to rice leaves. Compared to control plants treated with either *YFP*-dsRNA or water, effective protection was observed in plants treated with *Rs-DCTN1+SAC1*-dsRNA (Fig. 4a, b). Polygalacturonase (PG), a pectin degrading enzyme that hydrolyzes α-1,4 glycosidic bonds of pectic acid, is a known virulence factor of *R. solani* (Rao et al., 2019). Stable expression of a PG-RNAi construct in rice suppresses *R. solani* infection (Rao et al., 2019). We found that application of *Rs-PG*-dsRNA also exhibited a reduction in disease symptoms and biomass compared to water and *YFP*-dsRNA treated controls (Fig. 4a, b). Further, the mRNA expression levels of the corresponding genes were reduced in *R. solani* after the dsRNA treatment (Fig. 4c). Therefore, externally applied *Rs-DCTN1+SAC1*-dsRNA or *Rs-PG*-dsRNA protected the plants from *R. solani* infection by targeting and silencing *R. solani* genes.

**Figure 4.**
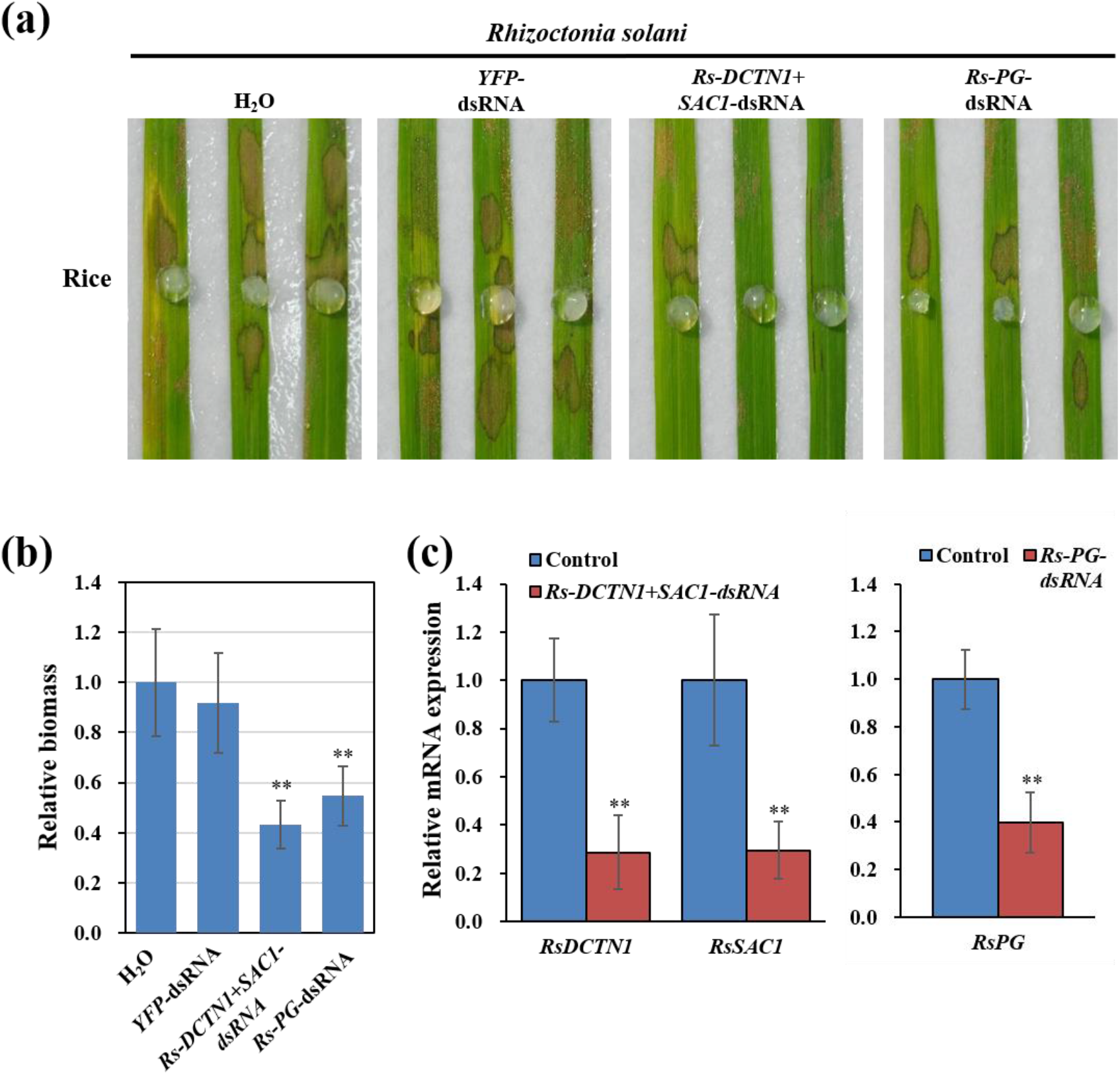
Topical application of pathogen gene-targeting dsRNAs inhibited the *R. solani* virulence. **(a)** Rice leaves were inoculated with *R. solani* mycelium plugs after treating with controls (water and *YFP*-dsRNA), *Rs-DCTN1+SAC1-*dsRNA or *Rs-PG*-dsRNA (40 ng μl^−1^). Pictures were taken at 3 dpi. **(b)** Relative biomass of *R. solani* was calculated by examining the expression of *Rs-Actin* by qRT-PCR, which is normalized to *Os18S rRNA*; error bars represent the SD of three replicates. Asterisks (**) indicate statistically significant differences (*P* < 0.01, Student’s t-test) between the treatment and the water control. **(c)** The mRNA expression levels of *RsDCTN1*, *RsSAC1* and *Rs-PG* were detected in *Rs-DCTN1+SAC1*-dsRNA and *Rs-PG*-treated *R. solani* by qRT-PCR analysis, respectively. Asterisks (**) indicate statistically significant differences (*P* < 0.01, Student’s t-test). Similar results were observed from three biological replicates.

### RNA is not stable soil and pretreatment of roots with dsRNA inhibited the infection of *V. dahliae*

*V. dahliae* is a root pathogen that causes wilt disease in hundreds of plant species. Previously, HIGS strategies have been successful to suppress *V. dahliae* infection (Song and Thomma, 2018; Wang et al., 2016; Xu et al., 2018). However, it is unknown whether direct application of dsRNAs can effectively control *V. dahliae* virulence. The *V. dahliae* genome encodes *DCTN1* and *SAC1* genes, but not the *VPS51* gene. As with the other pathogens, we generated *Vd-DCTN1+SAC1*dsRNA and *Vd-DCL1*+*DCL2* dsRNA. We found that dsRNA is not stable in the soil (Fig. 5a) and direct application of dsRNA to the soil didn’t protect the plant from *V. dahliae* infection. So we dipped Arabidopsis roots in *Vd-DCL1/2*-dsRNA or *Vd-DCTN1+SAC1*-dsRNA mixed with *V. dahliae* spores, then planted the seedlings in soil. These treated plants showed reduced disease symptoms in comparison to control plants treated with either *YFP-* dsRNA+spores, or only spores (Fig. 5b, c). Furthermore, *V. dahliae* treated with *Vd-DCL1/2*-dsRNA or *Vd- DCTN1+SAC1*-dsRNA showed a clear reduction in targeted mRNA expression levels (Fig. 5d). Although pretreatment of roots with dsRNA could protect the plants to a certain degree, it is not practical in the field. Therefore, innovative RNA protection strategies, such as using nanoparticles or special formula, are desired to protect the RNAs in the soil to increase the SIGS efficiency against soil-borne pathogens.

**Figure 5.**
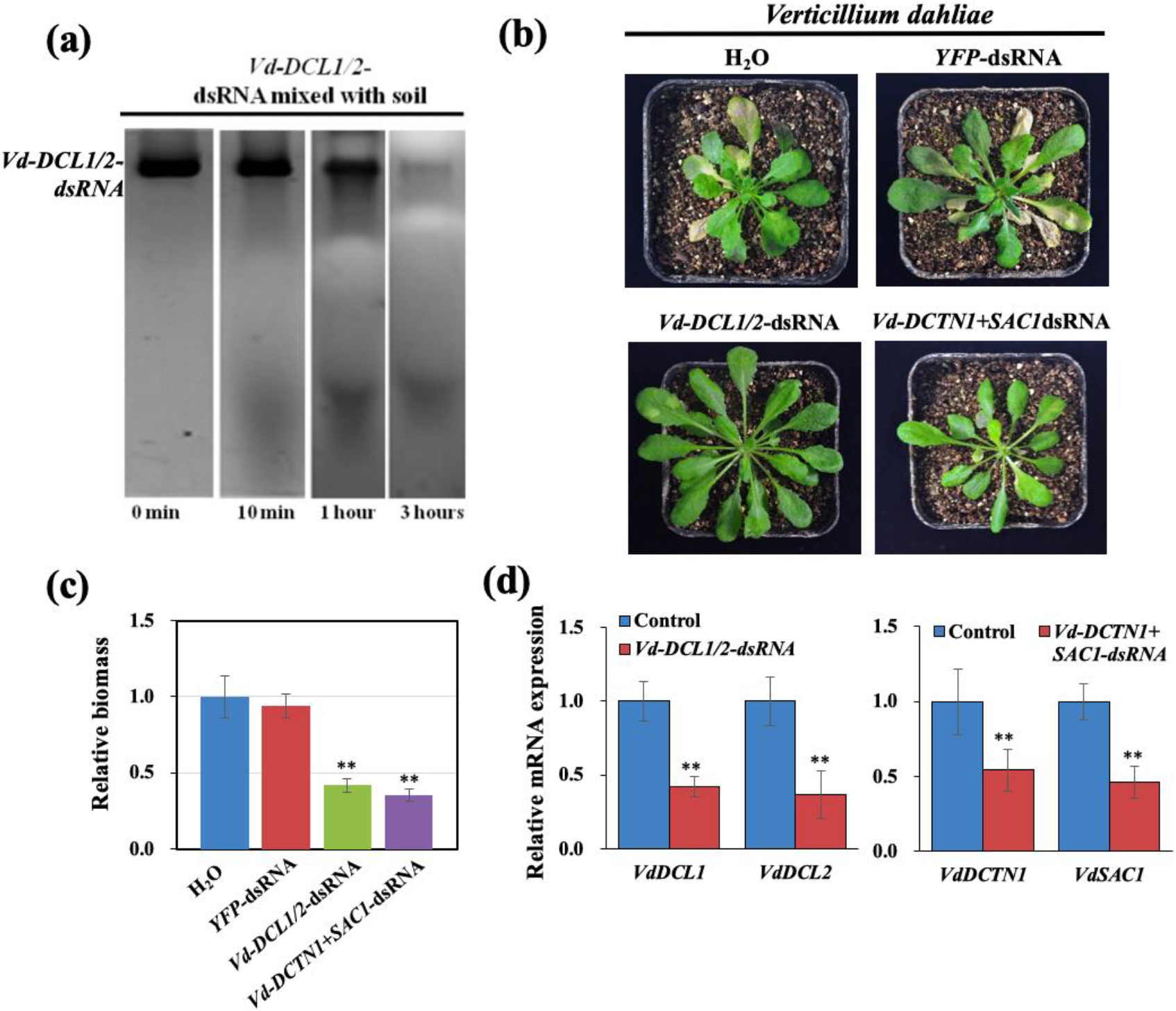
RNA is not stable in the soil and pretreatment of roots with pathogen gene-targeting dsRNA reduced the infection of *V. dahliae.* **(a)** Northern blot analysis of Vd-DCL1/2 dsRNAs was performed over a time course after mixed with soil. **(b)** *Arabidopsis* plants were uprooted and incubated in *V. dahliae* spore suspension (10^6^ spores/ml) with controls (water and *YFP*-dsRNA), *Vd-DCL1/2*-dsRNA, and *Vd-DCTN1+SAC1*-dsRNA (40 ng μl^−1^) treatments. Pictures were taken at 14 dpi. **(c)** Relative biomass of *V. dahliae* was calculated using by examining the expression of *Vd-actin* by qRT-PCR, which is normalized to *At-actin*; error bars represent the SD of three replicates. Asterisks (**) indicate statistically significant differences (*P* < 0.01, Student’s t-test) between the treatment and the water control. **(d)** The mRNA expression levels of *VdDCL1 and VdDCL2* were detected in *VdDCL1/2*-dsRNA-treated, as well as the mRNA expression levels of VdDCTN1 and VdSAC1were detected in VdDCTN1+VdSAC1-dsRNA-treated *V. dahliae* by qRT-PCR analysis. Asterisks (**) indicate statistically significant differences (*P* < 0.01, Student’s t-test) between the treatment and the water control. Similar results were observed from three biological replicates.

### Topical application of dsRNA cannot inhibit the virulence of *C. gloeosporioides*

We also assessed the influence of dsRNA on the virulence of *C. gloeosporioides*, a fungal pathogen that is unable to take up environmental dsRNA (Fig.1C). It was found that DCL proteins are important for the development and pathogenesis of *C. gloeosporioides* (Wang et al., 2017b). Here, we generated *Cg-VPS51+DCTN1+SAC1*-dsRNA and *Cg-DCL1/2-*dsRNA from the non-conserved regions of these genes. We then tested the efficacy of these dsRNAs in controlling *C. gloeosporioides* disease by separately applying the dsRNAs to the surface of various fruits (e.g., cherry, apple, and tomato). We found that all the fruits treated with pathogen gene targeting dsRNA displayed similar severity of disease symptoms as control fruits treated with water or *YFP*-dsRNA (Fig. S7a, b). Moreover, *C. gloeosporioides* treated with dsRNAs showed no reduction in the expression of targeted mRNAs in comparison to the controls (Fig. S7c), indicating that SIGS is not effective on controlling *C. gloeosporioides*.

### *P. infestans* takes up external dsRNA with low efficiency and topical application of dsRNA that targets *P. infestans* virulence genes fails to inhibit disease

In addition to fungi, oomycetes also cause severe crop diseases worldwide. For example, potato late blight, caused by *P. infestans*, induces the death of potato stems and leaves and leads to tuber rotting (Leesutthiphonchai et al., 2018; Shattock, 2002). We measured the dsRNA uptake efficiency of *P. infestans*. After incubating hyphae with fluorescein-labeled *YFP*-dsRNA and subsequently treating with MNase, weak fluorescence signals were detected in some *P. infestans* hyphae in plug-inoculated rye-sucrose medium cultures, whereas no signal was observed in hyphae derived from the germination of zoospores 12 h post-inoculation in rye medium (Fig. 6a and Fig. S8). Fluorescence signals were detected in some zoospore cysts and sporangia (Fig. 6b, c, and Fig. S8). However, fluorescence was only observed in approximately 5–10% of sporangia and zoospore cysts.

**Figure 6.**
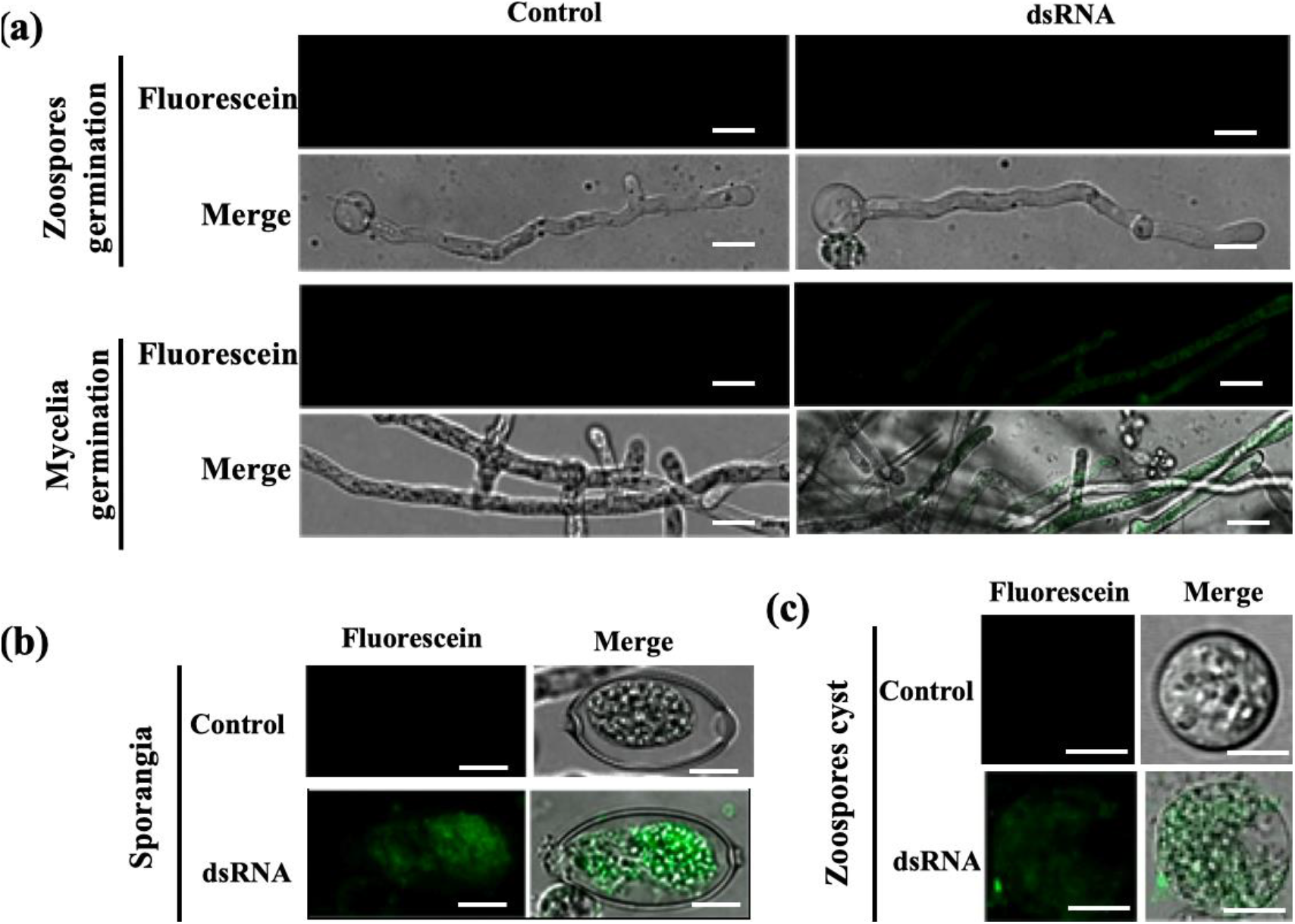
Different cell types of *P. infestans* take up external fluorescein-labeled *YFP*-dsRNA with different efficiencies. **(a)** Weak fluorescent signals were observed in *P. infestans* hyphae from plug-inoculated cultures treated with fluorescein-labeled *YFP*-dsRNA, whereas no signal was observed in germinated cysts (hyphae germinated from zoospores) treated with fluorescein-labeled *YFP*-dsRNA after culturing on rye agar medium for 10 h. Scale bars = 20 μm. (a-b) A weak fluorescence signal was observed in *P. infestans* sporangia **(b)** and zoospore cysts **(c)** treated with fluorescein-labeled *YFP*-dsRNA for 10 h after culturing on rye agar medium. Scale bars = 10 μm.

To examine whether SIGS inhibited *P. infestans* infection, we selected several target genes that are important for *P. infestans* virulence. Silencing of the *Pihmp1* gene that encodes the haustorium-specific membrane protein PiHMP1 has been shown to lead to a loss of pathogenicity (Avrova et al., 2008). Additionally, the expression of the *P. infestans* G-protein beta-subunit gene *PiGPB1* is required for sporangium formation (Latijnhouwers and Govers, 2003). Oomycetes possess small RNA biosynthesis pathways similar to those of other eukaryotes (Vetukuri et al., 2011), *PiDCL1* was also chosen as a target gene. The vesicle trafficking pathway genes *DCTN1* and *SAC1,* but not *VPS51* were identified in the *P. infestans* genome and were also selected as targets. dsRNAs of all target genes, *PiDCL1-*, *PiHMP1-*, *PiPGB1-*, and *DCTN1+SAC1*, were separately applied to potato leaves before inoculation with *P. infestans* zoospores. These dsRNA treatments did not reduce or slow disease development (Fig. S9a), as the size of disease lesions were indistinguishable from control-treated plants (Fig. S9a, b). Moreover, *P. infestans* treated with dsRNAs showed no significant reduction in targeted mRNA expression levels (Fig. S9c).

### Examine the longevity of dsRNA-mediated plant protection

We next investigated the protection effect on pre-harvesting crops over time by tracking the SIGS-RNAi efficiency against *B. cinerea* on tomato plants in a greenhouse assay. To simulate a commercial application, we treated the leaves of 5- to 6-week-old tomato plants by spraying 100 ng μl^−1^ *Bc-DCL1/2-* or *Bc-VPS51+DCTN1+SAC1*-dsRNA. We used higher concentration of dsRNA here because tomato leaves are more susceptible to *B. cinerea* and are infected more rapidly than fruits. Subsequently, the leaves were inoculated with *B. cinerea* spores at 1, 3, 7 and 14 dpt (Fig. 7a). Both *Bc-DCL1/2-* and *Bc-VPS51+DCTN1+SAC1-*dsRNA showed significant reduction in fungal growth (Student’s t-test at a<0.05) compared to the controls at 1-, 3- and 7-dpt, while at 14-dpt the effect was insignificant (Fig. 7b). Taken together, the dsRNAs remained effective on protecting the plants for up to 7 days, although the RNAi effect is reduced over time, likely due to dsRNA degradation in the environment.

**Figure 7.**
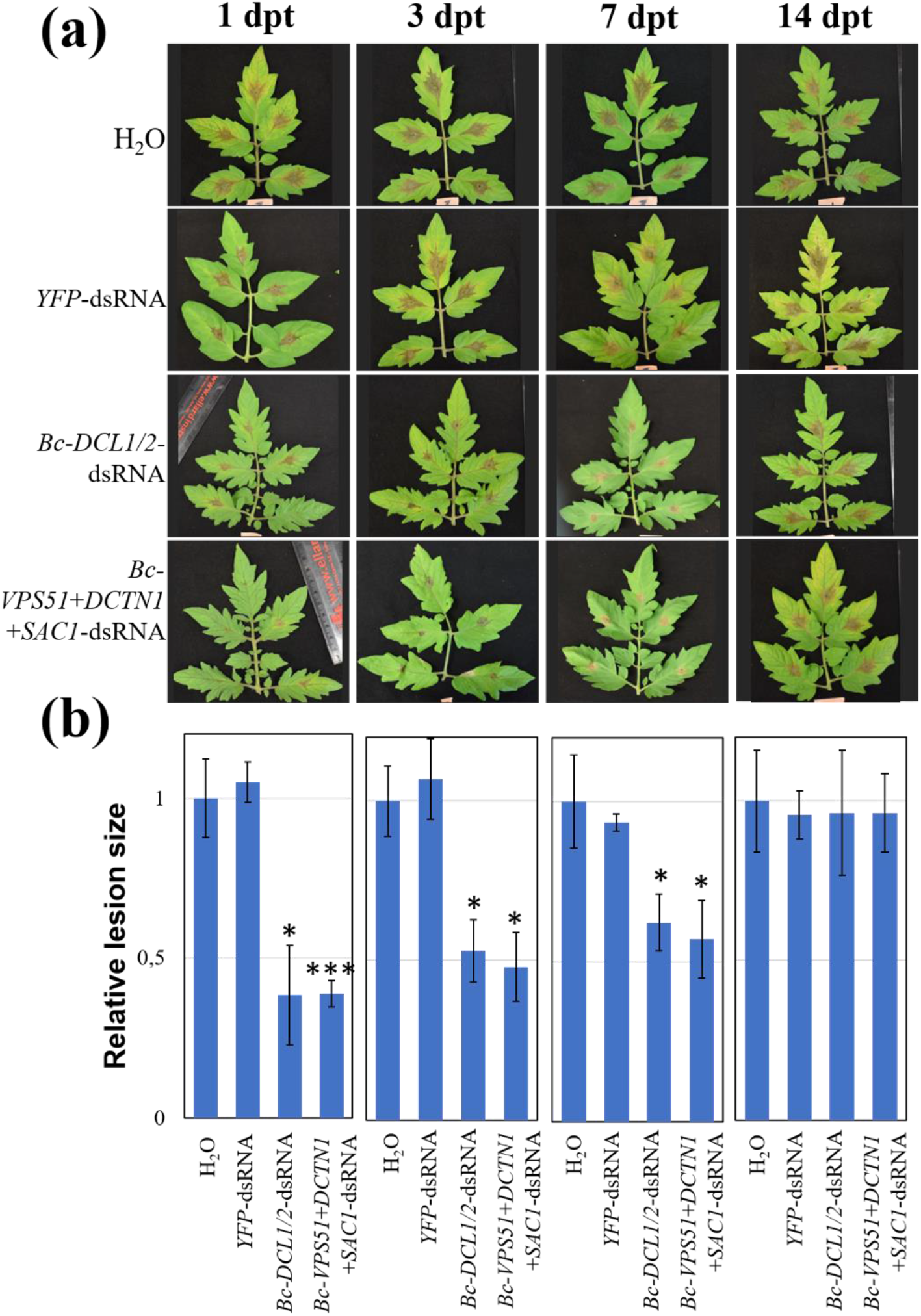
Examine the longevity of dsRNA-mediated plant protection. **(a)** Tomato leaves were inoculated with *B. cinerea* spores after spraying with controls (water or YFP-dsRNA) or *Bc-VPS51+DCTN1+SAC1-*dsRNA dsRNA or *Bc-DCL1/2*-dsRNA (100 ng μL^−^1) at 1, 3, 7 and 14 dpt. The relative lesion sizes were measured 4 dpi **(d)**. Error bars indicate the SD of at least three independent biological replicates with total of 45 leaflets in each replicate, and the statistical significance (Student’s t-test) by *, *P* < 0.05; ***, *P* < 0.001 between the treatment and the water control.

## Discussion

RNA-based disease management products offer an important eco-friendly alternative to standard chemical pesticides for controlling agricultural losses caused by pests and disease (Taning et al., 2020). SIGS has been shown to be effective on controlling gray mold disease caused by *B. cinerea* (Cai et al., 2018a; McLoughlin et al., 2018; Wang et al., 2016; Wang and Jin, 2017), *Fusarium* head blight caused by *F. graminearum* (Koch et al., 2016), and white mold caused by *S. sclerotiorum* (McLoughlin et al., 2018). To explore whether this RNA-based pathogen control strategy can be adapted across a multitude of eukaryotic pathogens, we investigated the dsRNA uptake efficiency of different species of fungi and oomycetes, and their pathogenicity after dsRNA treatment. Here, we provide solid evidence that the environmental dsRNA can be taken up by multiple fungal species with different uptake efficiencies, which determines the effectiveness of SIGS.

In the present study, we found that the uptake efficiency of RNA was significantly different among various fungi. Moreover, some fungi cannot take up dsRNA from the environment, such as *C. gloeosporioides*, thus limiting the application of SIGS in the control of these fungal diseases. Taken together, our results demonstrate that externally applied dsRNA can protect plants from fungal pathogens with high RNA uptake efficiency such as *B. cinerea, S. sclerotiorum*, *A. niger* and *R. solani*, but not against pathogens with low or no RNA uptake efficiency such as *C. gloeosporioides* and oomycete pathogen *P. infestans.* Thus, RNA uptake efficiency is crucial for controlling pathogens by SIGS. Recently, researchers have found that *Zymoseptoria tritici* was incapable of exogenous dsRNA uptake, which may limit RNAi approaches as a control measure of this fungal species also (Kettles et al., 2019). Further, we found that different cell types have different dsRNA uptake efficiency, such as *P. infestans,* its RNA uptake was observed in only a fraction of tissues. Thus, it was not surprising that treating plants with dsRNAs targeting essential *P. infestans* genes did not suppress *P. infestans* virulence. However, previous work has found HIGS strategies to be effective in preventing infection by the oomycete pathogen *Bremia lactucae* (Govindarajulu et al., 2015). and *P. infestans* (Jahan et al., 2015). This shows that *P. infestans* takes up plant-originated RNAs more efficiently, suggesting that mechanisms such as artificial vesicle-mediated delivery may increase RNA uptake efficiency in *P. infestans*. Notably, *P. infestans,* possesses a cell wall that is chemically dissimilar to those of fungi (Melida et al., 2013). This may also partially account for its less capable of taking up environmental RNAs.

The fungal cell wall is essential for viability, pathogenicity, and also regulates material exchange with the external environment (Cantu et al., 2009). The relative abundance and the chemical structures of the major components in the cell wall differ among different fungi (Bowman and Free, 2006; Diaz et al., 1992). Additionally, the number of chitinase genes in fungi display wide variations, from a single gene in *Schizosaccharomyces pombe*, to 36 genes in *T. virens* (Yang and Zhang, 2019). All these may impact the viscoelastic properties and dsRNA uptake efficiency. Walker *et al* have found that the cell wall of mammalian fungal pathogens *Candida albicans* and *Cryptococcus neoformans* allows for the uptake of liposome vesicles (Walker et al., 2018). These findings suggest that the fungal cell wall is deformable and viscoelastic to allow vesicles and large molecules, such as RNAs to pass through.

Finally, to better characterize the commercial potential of SIGS, we determined the longevity of plant protection against fungal infection conferred by dsRNAs. We found that naked dsRNAs can remain active up to a week but the efficacy of protection decreased over time. Innovative methods for enhancing RNA stability will be highly desirable for commercial use of SIGS. These methods include RNA modifications as well as the use of protective nanoparticles. An efficient carrier for SIGS-mediated plant protection is BioClay, which has been shown to sustain plant protection against viruses for weeks (Mitter et al., 2017). In applications against the insect pest, rice striped stem borer, researchers have found that carbon quantum dots (CQD) are efficient dsRNA carriers which can induce systemic RNAi in the insect target (Wang et al., 2020). Another class of nanoparticles worth exploring are artificial vesicles or liposomes, which are used in clinical contexts to deliver therapeutic RNAs (Bochicchio et al., 2014), and may mimic the natural pathway of extracellular-vesicle mediated sRNA transport from plants to pathogens. They may also have the potential to improve pathogen RNA uptake efficiency, ultimately leading to longer lasting and more robust plant protection against infections.

To conclude, SIGS disease management strategies show promising results in many, but not all, eukaryotic pathogens. Efficacy of SIGS is directly related to RNA uptake efficiency by the pathogen. Further optimization of delivery vectors and methods could improve the uptake efficiency in pathogens. Finally, before widespread commercial use of RNA-based disease management strategies is practical, it is crucial that environmental RNA-stability is enhanced.

## Methods

### Fungi and oomycete culture and infection conditions

*S. sclerotiorum* 1980, *R. solani* AG1-IA, *A. niger* CBS 513.88, *T. virens* Gv29-8, *V. dahliae* JR2, and *C. gloeosporioides* Nara gc5 were routinely cultured on PDA medium (24 g of potato dextrose broth and 15 g of agar per liter) at 25 °C. *B. cinerea* B05.10 was cultured on malt extract agar medium (20 g of malt extract, 10 g of Bacto protease peptone, and 15 g of agar per liter). Fungal mycelia used for genomic DNA and total RNA extraction was harvested from cultures grown on medium covered by a sterile cellophane membrane. Vegetables (lettuce and collard greens), rose petals, and fruits (tomato, cherry, grape and apple) purchased from local supermarkets were used as experimental host plant materials. For *B. cinerea* infection, the fungal spores were diluted in 1% sabouraud maltose broth buffer to a final concentration of 10^5^ spores/ml for drop inoculation of plants. For *S. sclerotiorum* infection, mycelium plugs (5 mm in diameter) were obtained from the expanding margins of colonies to inoculate vegetable leaves. Infected leaf tissues were cultured in a light incubator at 25 °C for 72 h and fruits for 120 h at constant and high humidity. For *R. solani* inoculations, *R. solani* was cultured for 48 h on PDA medium. A mycelium plug (5 mm diameter) of actively growing *R. solani* was placed onto the surface of rice leaves. The *V. dahliae* soil-inoculation assay was performed as previously described (Ellendorff et al., 2009). Fungal biomass quantification was performed as described previously (Gachon and Saindrenan, 2004). *C. gloeosporioides* inoculations were based on the standard protocol, with minor modifications (Seo et al., 2014). Fungal inoculation was conducted by applying 10 μl droplets of spore suspension (1 × 10^5^ spores per ml in distilled water) onto fruits. The inoculated fruits were placed in high humidity and dark conditions for 1 day to stimulate infection. For *A. niger* inoculations, the fungal spores were adjusted to 10^6^ spores/ml for drop inoculation of fruits. *P. infestans* 1306 culture and infection conditions were based on standard protocols (Ah-Fong et al., 2017). The strain was maintained in the dark in rye-sucrose agar at 18 °C. Developmental stages were obtained as described from rye media cultures. Potato plants used for infections were grown with a 12 h light/dark cycle (25 °C day, 350 μmol m^−2^ s^−1^ fluorescent light; 18 °C night) for 4–5 weeks, and then leaves were detached, placed on 0.8% water agar, inoculated with zoospores, and incubated at 16 °C with a 12 h light/dark cycle with 115 μmol m^−2^ s^−1^ illumination in a clear plastic bag to maintain high humidity. The lesion sizes of pathogen-infected plant materials were measured and calculated using ImageJ software.

### RNA extraction and qRT-PCR

RNA was extracted using the Trizol extraction method. Purified RNA was treated with DNase I (Thermofisher Scientific), then first-strand cDNA was synthesized using the SuperscriptTM III First-Strand Synthesis System (Thermofisher Scientific). RT-qPCR was performed using the CFX384 (Bio-Rad) Connect system using SYBR Green mix (Bio-Rad) with the following program: 95 °C for 15 min, followed by 40 cycles of 94 °C for 30 s, 50 °C for 30 s, and 72 °C for 30 s. Melt curves were generated to evaluate the fidelity of amplification. Expression levels were calculated using the ΔΔCt method.

### *S. sclerotiorum* protoplast transformation

*S. sclerotiorum* protoplast transformation was based on standard protocols with minor modifications (Rollins, 2003). Protoplasts were suspended at a concentration of 1 × 10^8^ protoplasts per ml in four parts STC (0.8 M sorbitol, 50 mM Tris-HCl, pH 8.0, and 50 mM CaCl_2_) and one-part SPTC media (0.8 M sorbitol, 40% polyethylene glycol 3500, 50 mM Tris-HCl, pH 8.0, and 50 mM CaCl_2_). A mixture of 5 μg of transforming DNA, 5 μl of spermidine (50 mM stock), and 5 μl of heparin (5 mg ml^−1^ in STC) was added to 100 μl of the protoplast suspension. The above mixtures were incubated on ice for 30 min, then gently mixed with 1 ml of SPTC solution and incubated at room temperature for 20 min. The transformation mixture was added to a 50 ml flask, then 10–20 ml of regeneration medium (RM; 274 g of sucrose, 1 g of yeast extract, and 1 g of casamino acid per liter) was added. The mixture was shaken at 100 rpm and left overnight at 25 °C. The protoplasts were gently mixed in 200 ml of regeneration medium RMA (RM containing 15 g agar per liter) at 43 °C, poured into 9 cm diameter Petri dishes (10 ml per dish), and incubated at 25 °C. After 24 h, the plates were overlaid with 10 ml of selective medium containing 100 μg ml^−1^ hygromycin B. Transformants were obtained 4–10 days after transformation and transferred to fresh plates of PDA containing 100 μg ml^−1^ of hygromycin B. Hygromycin-resistant transformants were hyphal tip transferred a minimum of three times on PDA containing 100 μg of hygromycin per ml. Primers used for *S. sclerotiorum* knock-down mutant construction and detection are listed in Table S1.

### *In vitro* synthesis of dsRNA

*In vitro* synthesis of dsRNA was based on established protocols (Wang *et al.*, 2016). Following the MEGAscript® RNAi Kit instructions (Life Technologies, Carlsbad, CA), the T7 promoter sequence was introduced at both 5’ and 3’ ends of the RNAi fragments by PCR. After purification, the DNA fragments containing T7 promoters at both ends were used for *in vitro* transcription. Primers used for *in vitro* synthesis of dsRNAs are listed in Table S1, and the sequences of dsRNAs are listed in Table S2.

### Fungal uptake of fluorescein-labeled dsRNA *in vitro*

*In vitro* synthesis of fluorescein-labeled dsRNA was based on established protocols (Wang *et al.*, 2016). *YFP*-dsRNA was labeled using the fluorescein RNA Labeling Mix Kit following the manufacturer’s instructions (Sigma, St. Louis, MO). For confocal microscopy examination of fluorescent dsRNA uptake by fungal mycelium, 5 μl of 150 ng μl^−1^ fluorescent dsRNA was applied to 10 μl of 10^5^ spores ml^−1^ of *B. cinerea*, *V. dahliae*, *A.niger*, *T. virens*, *P. infestans*, and *C. gloeosporioides.* For *B. cinerea*, *A. niger*, *P. infestans*, the dsRNA-spore mixture was grown on PDA medium applied directly to the microscope slides surface up to 12 h. For *V. dahliae* and *T. virens*, the incubation period was extended up to 24-30 h since spore germination is slower in these species. *S. sclerotiorum* and *R. solani* mycelium plugs (5 mm in diameter) were placed on slides containing PDA medium and 20 μl of 150 ng μl^−1^ fluorescent dsRNA was added to the fungi, and incubated for 10–12 h before confocal imaging. All fungal growth slides were cultured in darkness at 25 °C. Germinating spores or mycelium was treated with KCl buffer (control) or 75 U micrococcal nuclease enzyme to degrade the dsRNA in the surface of spores or mycelium (dissolved in KCl buffer) before observation at 37 °C for 30 min. The fluorescent signal was analyzed using a Leica SP5 confocal microscope.

### External application of dsRNA on the surface of plant materials

For dsRNAs applied to *S. sclerotiorum*, *C. gloeosporioides* and *P. infestans*, the dsRNAs were adjusted to a concentration of 40 ng μl^−1^ with RNase-free water before use. For *in vitro* synthetic *YFP*-dsRNA, *Ss-DCL1/2*, and *Ss-VPS51+DCTN1+SAC1*-dsRNA, 20 μl of dsRNA (40 ng μl^−1^) was dropped onto the surface of each plant sample. Then, 5 mm diameter *S. sclerotiorum* mycelium plugs were applied to the same site as the dsRNA treated area. A total of 20 μl dsRNA (40 ng μl^−1^) was dropped onto the surface of each fruit sample and then *C. gloeosporioides* spores (1 × 10^5^ per ml) were applied to the same dsRNA-treated area. For *in vitro* synthetic *YFP*-, *PiDCL1-*, *PiHMP1-*, and *PiPGB1*-dsRNA, 20 μl of dsRNA (40 ng μl^−1^) was dropped onto the surface of potato leaves, then *P. infestans* zoospores were applied to the same the same dsRNA treated area. For dsRNAs that were applied in controlling *B. cinerea* and *A. niger,* the dsRNAs were adjusted to a concentration of 20 ng μl^−1^ for the fruits, vegetables and petals. For applications on tomato leaves, dsRNA (100 ng μl^−1^) was sprayed onto the leaf surface. In all the above experiments, fungal inoculation was performed after dsRNA treatment as described in Wang *et al.*, 2016. For the time-course experiments, fungal inoculation was performed at the days post RNA treatment indicated in figure legends. For dsRNAs applied to control *V. dahliae*, *Arabidopsis* roots were dipped in a mixture of *V. dahliae* spores and the dsRNA (20 ng μl^−1^) for 3 min (Ellendorff *et al.*, 2009), then seedlings were planted in soil.

## Acknowledgments

This work was supported by grants from National Institute of Health (R01 GM093008 and R35 GM136379-01), National Science Foundation (IOS-1557812 and IOS-2017314) and the CIFAR ‘Fungal Kingdom’ fellowship to H.J.; by the Jiangsu Agricultural Science and Technology Innovation Fund of China (CX(19)3103) to D.N.; and two graduate student fellowships, one is from National Science Foundation (Research Traineeship grant DBI-1922642) to R.H. in H.J.’s lab, and the other is from AMPELOS Grape nurseries organization, Italy to L.C. in B.M.’s lab. We thank Suomeng Dong for providing the pathogen *P. infestans* and Shimei Wang for providing the pathogen *Aspergillus niger.* This material by m-CAFEs Microbial Community Analysis & Functional Evaluation in Soils, a Project led by Lawrence Berkeley National Laboratory is based upon work supported by the U.S. Department of Energy, Office of Science, Office of Biological & Environmental Research under contract number DE-AC02-05CH11231.

## Author contributions

H.J. conceived the idea. L.Q., C.L., L.C., A.A-F., J.H., H.S.J, R.H., J.N.S. and D.N. performed the experiments. L.Q., L.C. and H.J. drafted the manuscript. H.J., D.N., A.A-F., J.H., H.Z., N.L.G., H.S.J,B.M., R.H. and J.N.S. revised the manuscript. H.J. and D.N. designed the experiments and supervised the study. All authors read and approved of its content.

## Supplementary information

The following Supporting Information is available for this article:

**Figure S1.** Colony morphology of different types of fungi.

**Figure S2.** Mycelium morphology of different types of fungi.

**Figure S3.** Multiple fungal dsRNA uptake efficiencies were observed over a time course.

**Figure S4.** Fluorescein-labeled *YFP*-dsRNA was observed in *S. sclerotiorum* protoplasts after MNase treatment.

**Figure S5.** Accumulation of fluorescein signal intensities in different fungal cells.

**Figure S6.** *SsVPS51*, *SsDCTN1*, and *SsSAC1* knockout mutants displayed significantly reduced virulence compared to the wild-type strain.

**Figure S7.** Topical application of pathogen gene-targeting dsRNAs had no effect on *C. gloeosporioides* infection and disease.

**Figure S8.** Accumulation of fluorescein signal intensities in different *P. infestans* cells.

**Figure S9.** *YFP-*, *PiHMP1-*, *PiPGB1*-, *PiDCL1-*, and *Pi-DCTN1+SAC1-*dsRNAs did not inhibit *P. infestans* infection on potato leaves.

**Table S1.** Primers used in the study

**Table S2.** The sequences of dsRNAs in the study.

