## supplemental data for "Spray-induced gene silencing for disease control is dependent on the efficiency of pathogen RNA uptake"

**This file includes:**

**Figures S1-S9 and Table S1-S2**

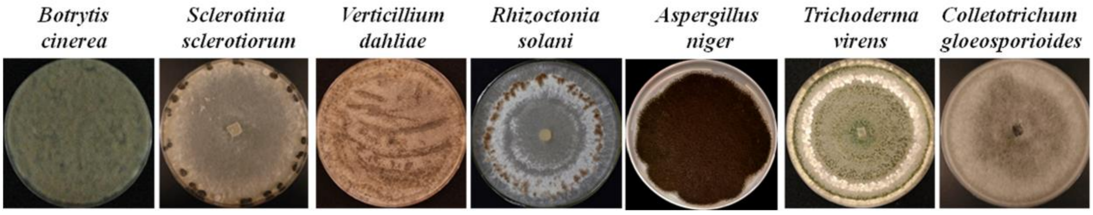


**Figure S1. Colony morphology of different types of fungi.** Colony morphology pictures of *B. cinerea*, *S. sclerotiorum*, *V. dahliae*, *R. solani*, *A. niger*, *T. virens,* and *C. gloeosporioides*.

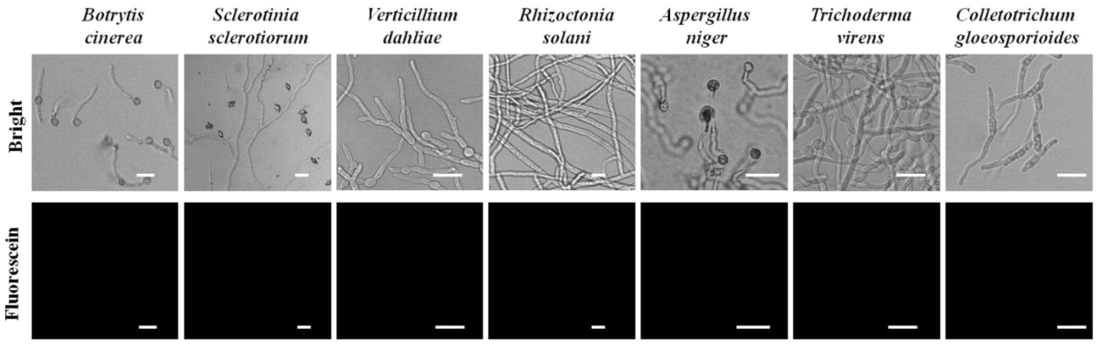


**Figure S2. Mycelium morphology of different types of fungi**. Mycelium morphology pictures of *B. cinerea*, *S. sclerotiorum*, *V. dahliae*, *R. solani*, *A. niger*, *T. virens*, and *C. gloeosporioides* were taken by confocal, all fungal cells show no fluorescent signals. Scale bars = 20 μm.

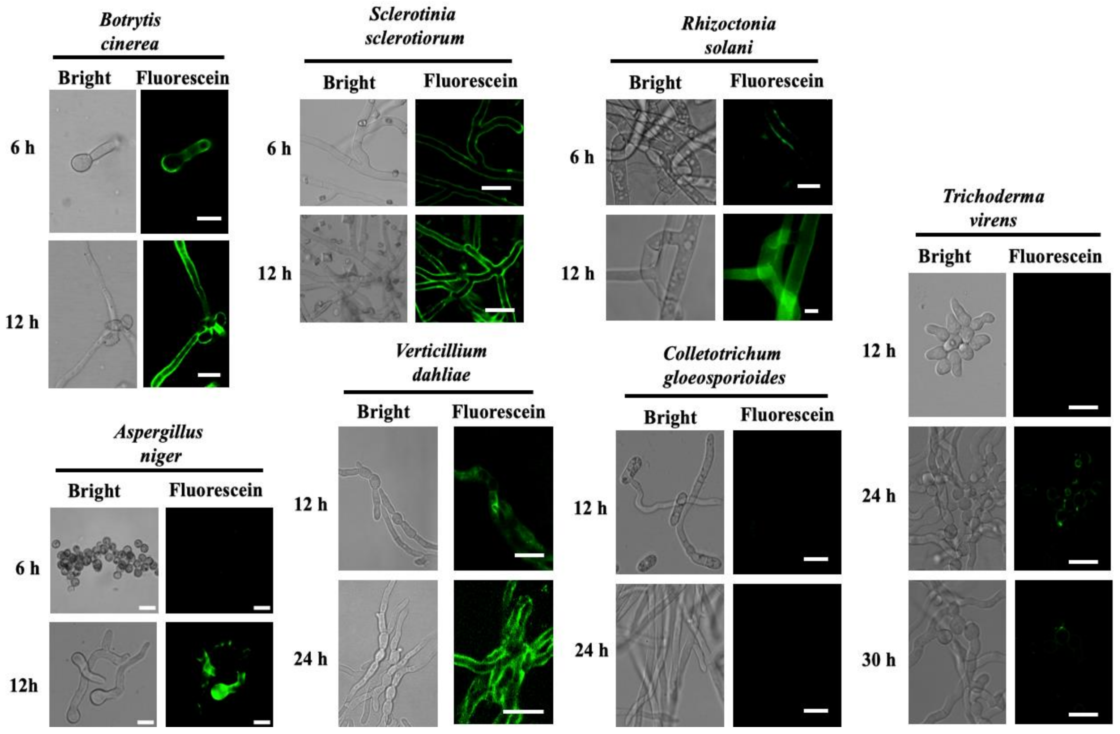


**Figure S3. Multiple fungal dsRNA uptake efficiencies were observed over a time course.** Fluorescein-labeled *YFP*-dsRNA was added to *S. sclerotiorum*, *R. solani*, *A. niger*, *C. gloeosporioides*, *V. dahliae*, *B. cinerea* and *T. virens* hyphae, as well as spores, and the fluorescence signals were observed over time courses uses a Leica confocal microscope. Micrococcal nuclease (MNase) treatment was performed 30 min before image acquisition. Scale bars = 20 µm.

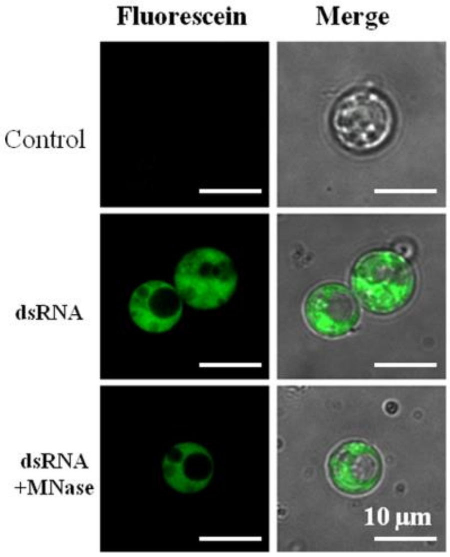


**Figure S4. Fluorescein-labeled *YFP*-dsRNA was observed in *S. sclerotiorum* protoplasts after MNase treatment.** Fluorescein-labeled *YFP*-dsRNA was applied to germinated *S. sclerotiorum* mycelium and the protoplasts were isolated after culturing for 48 h. The fluorescent signals were detected within fungal protoplasts and MNase enzyme treatment did not reduce the fluorescent signal intensities. Similar results were obtained over three biological replicates. Scale bars = 10 µm.

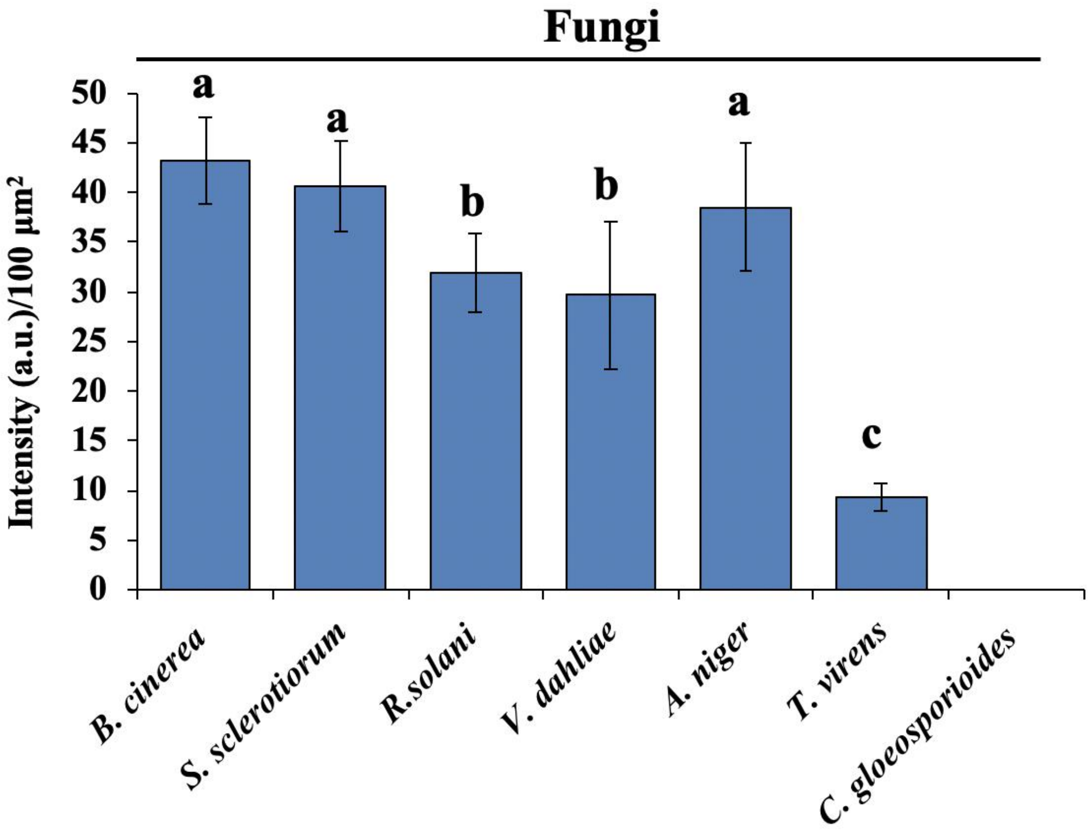


**Figure S5. Accumulation of fluorescein signal intensities vary in different fungal cells.**

The fluorescein signal intensities were quantified using ImageJ software, which could compare the fluorescence intensity of the same area between different cells. The letters above the bars indicate significant difference at a value of $P < 0.05$ (Duncan's multiple range test). Similar results are obtained from three biological repeats.

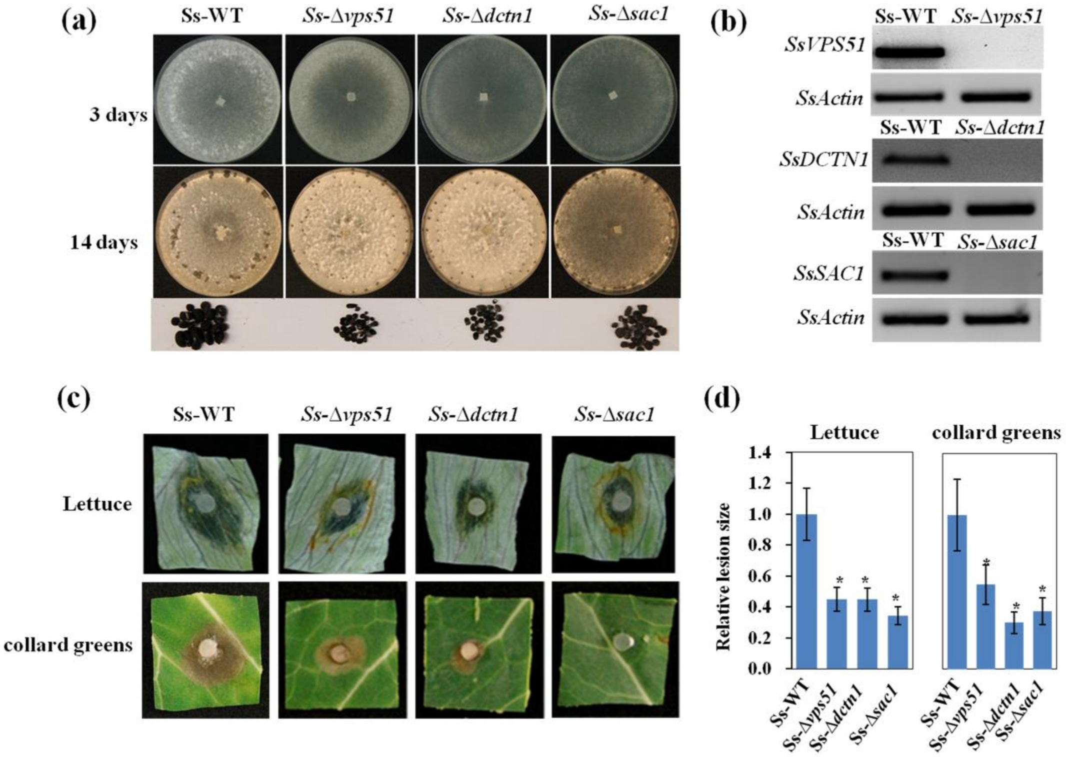


**Figure S6. *SsVPS51*, *SsDCTN1*, and *SsSAC1* knockout mutants displayed significantly reduced virulence compared to the wild-type strain.**

**(a)** Phenotypes of the *SsVPS51*, *SsDCTN1*, and *SsSAC1* knockout mutant strains grown on PDA solid medium at 25 °C. Photographs were taken after 3 and 14 days of fungal growth. **(b)** The expression levels of each gene in the knockout mutant strain were measured by RT-PCR. The *S. sclerotiorum Actin* gene was used as the internal expression control. Similar results were observed for the three biological replicates. **(c)** *Ss-Δvps51*, *Ss-Δdcnt1,* and *Ss-Δsac1* strains showed reduced virulence on lettuce and collard greens compared with Ss-WT. **(d)** The relative lesion sizes were measured at 3 dpi for lettuce and collard greens using ImageJ and error bars represent the SD of 10 plant samples. Asterisks (*) indicate statistically significant differences (P < 0.05, Student's t-test). Similar results were observed for the three biological replicates.

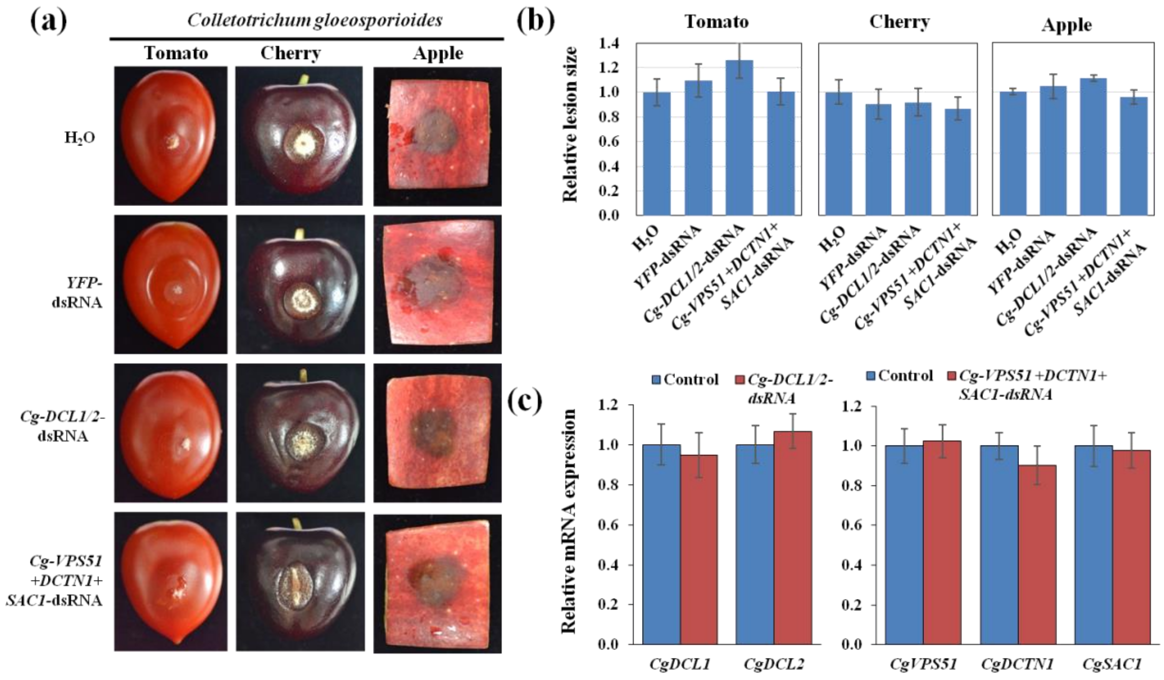


**Figure S7. Topical application of pathogen gene-targeting dsRNAs had no effect on *C. gloeosporioides* infection and disease.**

**(a)** Tomatoes, cherries, and apples were inoculated with *C. gloeosporioides* spores after treating with controls (water and *YFP*-dsRNA), *Cg-DCL1/2*-dsRNA or *Cg-VPS51+DCTN1+SAC1*-dsRNA (20 µl of 40 ng $\mu l^{-1}$). Pictures were taken at 5 dpi. **(b)** The relative lesion sizes were measured at 5 dpi using ImageJ software. Error bars indicate the SD of 10 samples. **(c)** The mRNA expression levels of *CgDCL1* and *CgDCL2* were detected in *Cg-DCL1/2*-dsRNA-treated *C. gloeosporioides*, and *CgVPS51*, *CgDCTN1* and *CgSAC1* were detected in *Cg-VPS51+DCTN1+SAC1*-dsRNA-treated *C. gloeosporioides*. Error bars indicate the SD of three technical replicates. Similar results were observed from three biological replicates.

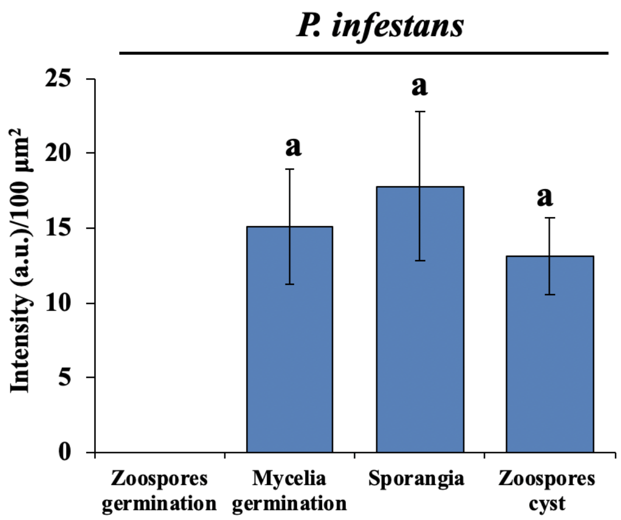


**Figure S8. Accumulation of fluorescein signal intensities in different *P. infestans* cells.** The fluorescein signal intensities were quantified using ImageJ software, which could compare the fluorescence intensity of the same area between different cells. The letters above the bars indicate significant difference at a value of $P < 0.05$ (Duncan's multiple range test). Similar results are obtained from three biological repeats.

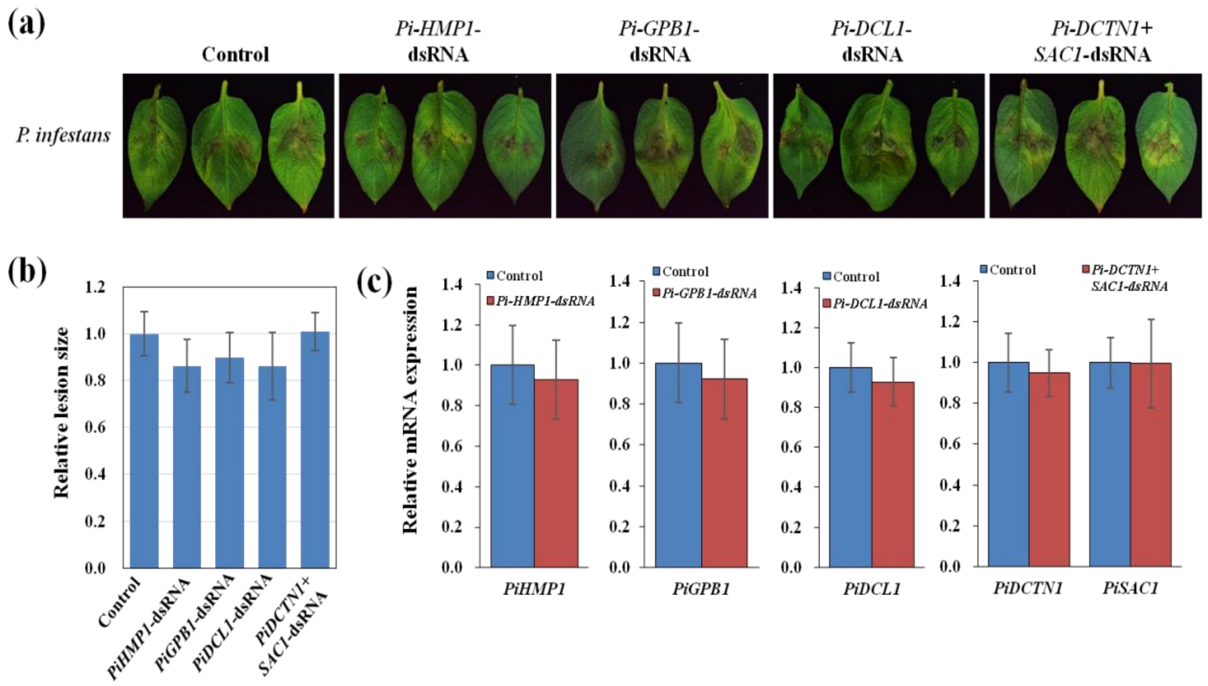


**Figure S9.** ***YFP-*****,** ***PiHMP1-*****,** ***PiPGB1-*****,** ***PiDCL1-*****, and** ***Pi-DCTN1+SAC1*****-dsRNAs did not inhibit** ***P. infestans*** **infection on potato leaves.**

**(a)** Potato leaves were treated with *YFP-*, *PiHMP1-*, *PiPGB1-*, *PiDCL1-*, and *Pi-DCTN1+SAC1*-dsRNAs (40 ng μl$^{-1}$) and then inoculated with *P. infestans*, imaged at 3 dpi. **(b)** Relative lesion sizes were measured at 3 dpi using ImageJ software. Error bars indicate the SD of 10 samples. Similar results were observed in the three biological replicates. **(c)** The mRNA expression levels of *PiHMP1*, *PiPGB1*, *PiDCL1*, *PiDCTN1* and *PiSAC1* were detected in corresponding dsRNA-treated *P. infestans* by qRT-PCR analysis. Similar results were observed from three biological replicates.

**Table S1.** Primers used in the study

| Primer Name | Primer Sequence (5′-3′) | Primer Description |
|---|---|---|
| *YFP*-dsRNA-T7-F | TAATACGACTCACTATAGGGAGA ATGGTGAGCAAGGGCGAGGA | Template DNA for *in vitro* *YFP*-dsRNA synthesis |
| *YFP*-dsRNA-T7-R | TAATACGACTCACTATAGGGAGA TTACTTGTACAGCTCGTCCATGC | |
| *SsDCL1*-dsRNA-T7-F | TAATACGACTCACTATAGGGAGA CATTTCGGAGGCGGTCG | Template DNA for *in vitro* *Ss-DCL1/2* synthesis |
| *SsDCL1*-dsRNA-T7-R | GATTATGTTCTGCCTCTTGCCAT GCCTTGGACATATTGAACAG | |
| *SsDCL2*-dsRNA-T7-F | CTGTTCAATATGTCCAAGGCATG GCAAGAGGCAGAACATAATC | |
| *SsDCL2*-dsRNA-T7-R | TAATACGACTCACTATAGGGAGA CGTAATATCATGTCCTCGG | |
| *SsVPS51*- dsRNA-T7-F | TAATACGACTCACTATAGGGAGA GCATTGGCTGCGCTGAAGAC | Template DNA for *In vitro* *Ss-VPS51+DCTN1+SAC1* synthesis |
| *SsVPS51*- dsRNA-T7-R | CCGTCAATCTCTGGCATTGACCT TGAAGAGTCTTTGTGAATC | |
| *SsDCTN1*-dsRNA-T7-F | GATTCACAAAGACTCTTCAAGGT CAATGCCAGAGATTGACGG | |
| *SsDCTN1*-dsRNA-T7-R | CCTGCTTGGAAATGTAAGCGTTT GGGTGATAGTTTGAGTC | |
| *SsSAC1*-dsRNA-T7-F | GACTCAAACTATCACCCAAACGC TTACATTTCCAAGCAGG | |
| *SsSAC1*-dsRNA-T7-R | TAATACGACTCACTATAGGGAGA CTGATAATGGCGATGCTATC | |
| *SsVPS51*-KO-5-F | GGGGTACCGGACGAATGGGAGT TATG | Construct *SsVPS5*1 ko mutant |
| *SsVPS51*-KO-5-R | CCCTCGAGCTCTTCGCCAACAAC CG | |
| *SsVPS51*-KO-3-F | AACTGCAGGTACATCTCGATTCC CAAAT | |
| *SsVPS51*-KO-3-R | CGGGATCCCTGTTAAGATATCTT AAATTTC | |
| *SsDCTN1*-KO-5-F | GGGGTACCCGTACCTTCCATTTT CCTC | Construct *SsDCTN1* ko mutant |
| *SsDCTN1*-KO-5-R | CCCTCGAGGTGTATACATGTTGC CTGC | |
| *SsDCTN1*-KO-3-F | AACTGCAGGTGTCAGCAATTCTA C | |
| *SsDCTN1*-KO-3-R | CGGGATCCCGGTTAACCAACATA GCC | |
| *SsSAC1*-KO-5-F | GGGGTACCCGTGCTCTTCTTGGC CTC | Construct *SsSAC1* ko mutant |
| *SsSAC1*-KO-5-R | CCCTCGAGCGGATGATGGTGAA GAATTTATTG | |
| *SsSAC1*-KO-3-F | CCCAAGCTTGGCAGCATGCTCTC AGC | |
| *SsSAC1*-KO-3-R | GGAATTCGTCGGACCAGCATCTT ATG | |
| *SsVPS51*-GT-5-F | CTTAGAGACTTGAGGAAGTC | *Ssvps51* ko mutant genotype identify |
| *SsVPS51*-GT-3-R | CTACTTCTATTGCTATTC | |
| *SsDCTN1*-GT-5-F | GATGAAGGAGTTCCCTTC | *Ssdctn1* ko mutant genotype identify |
| *SsDCTN1*-GT-3-R | CAGTACTAGATAATTCTAC | |

| | | |
|---|---|---|
| *SsSAC1*-GT-5-F | CTTGGTGCCCTAGCATATC | *Sssac1* ko mutant genotype identify |
| *SsSAC1*-GT-3-R | CAACGAAGATCACGTATAAAG | |
| *Ss-actin*-qRT-F | GAGCTGTTTTCCCTTCCATTGTC | qRT-PCR Gene Expression |
| *Ss-actin*-qRT-R | GACGACACCGTGCTCGATTGG | |
| *SsVPS51*-qRT-F | CTGGTACAGGCTCTTTGC | qRT-PCR *SsVPS51* expression |
| *SsVPS51*-qRT-R | CACCTGATGTCTCCTGGTTCG | |
| *SsDCTN1*-qRT-F | GGATATGCCACCAGCAGGC | qRT-PCR *SsDCTN1* expression |
| *SsDCTN1*-qRT-R | CACACTGCCGACAAATCTG | |
| *SsSAC1*-qRT-F | GCTCGGGACACATTGAGAAG | qRT-PCR *SsSAC1* expression |
| *SsSAC1*-qRT-R | GCCACAGGTTCATCATTCTACC | |
| *BcDCL1*-dsRNA-T7-F | TAATACGACTCACTATAGGGAGATGCGGAAGAACTTGAAGGTTTGCTACA | Template DNA for *In vitro Bc-DCL1/2* synthesis |
| *BcDCL1*-dsRNA-T7-R | GCAGCAAATGGCATCCGTCCAGATCTGGTCAACACACCAAG | |
| *BcDCL2*-dsRNA-T7-F | CTTGGTGTGTTGACCAGATCTGGACGGATGCCATTTGCTGC | |
| *BcDCL2*-dsRNA-T7-R | TAATACGACTCACTATAGGGAGAACTCTTGAGTACTTTCGCCAGCTCAC | |
| *BcVPS51*-dsRNA-T7-F | TAATACGACTCACTATAGGGAGATTCGTTCCAGGAGTTACACG | Template DNA for *In vitro Bc-VPS51+DCTN1+SAC1* synthesis |
| *BcVPS51*-dsRNA-T7-R | CTTTCGACGAGAGCACGAATGATGAGACAAGTGAGAGTCCA | |
| *BcDCTN1*-dsRNA-T7-F | TGGACTCTCACTTGTCTCATCATTCGTGCTCTCGTCGAAAG | |
| *BcDCTN1*-dsRNA-T7-R | CACTGCACTTTGAACAACGTCCAGCTTACAACTGTGCTCT | |
| *BcSAC1*-dsRNA-T7-F | AGAGCACAGTTGTAAGCTGGACGTTGTTCAAAGTGCAGTG | |
| *BcSAC1*-dsRNA-T7-R | TAATACGACTCACTATAGGGAGACCTTCAATGCTGCTGTAGAAG | |
| *Bc-actin*-qRT-F | TGCTCCAGAAGCTTTGTTCCAA | qRT-PCR Gene Expression |
| *Bc-actin*-qRT-R | TCGGAGATACCTGGGTACATAG | |
| *BcDCL1*-qRT-F | ACAATCCTATCTTTCGGAAGC | qRT-PCR *BcDCL1* Expression |
| *BcDCL1*-qRT-R | AGACTCTTCTTCTTGAAGACAG | |
| *BcDCL2*-qRT-F | GATTGTGCAAAGTCTCAACA | qRT-PCR *BcDCL2* Expression |
| *BcDCL2*-qRT-R | ATTGGGTTTGACTATATGTCTTA | |
| *BcVPS51*-qRT-F | GTTAGGCAAAGTAGGAGAAGG | qRT-PCR *BcVPS51* Expression |
| *BcVPS51*-qRT-R | CATCCATTCGCCCCAATACC | |
| *BcDCTN1*-qRT-F | GCTTCCCTTACCTCGAATTC | qRT-PCR *BcDCTN1* Expression |
| *BcDCTN1*-qRT-R | CAACCTCTAGTTCTTCCGGTC | |
| *BcSAC1*-qRT-F | CGAAAAGTTTCGAGGCATTTG | qRT-PCR *BcSAC1* Expression |
| *BcSAC1*-qRT-R | GAACTTAATGGGGTCAAAGCC | |
| *CgDCL1*-dsRNA-T7-F | TAATACGACTCACTATAGGGAGAGTCAGATGTCTATCCCGTTTCCTTG | Template DNA for *in vitro Cg-DCL1/2* synthesis |
| *CgDCL1*-dsRNA-T7-R | CTCGATGGGAAGCTGATCACCGACTTCAGAAAGCTTTGCCG | |

| | | |
|---|---|---|
| *CgDCL2*-dsRNA-T7-F | CGGCAAAGCTTTCTGAAGTCGGTGATCAGCTTCCCATCGAG | |
| *CgDCL2*-dsRNA-T7-R | TAATACGACTCACTATAGGGAGAGAAGCAGGTGAGGTCGTGTGC | |
| *CgVPS51*-dsRNA-T7-F | TAATACGACTCACTATAGGGAGAGTCTGGTCGTGCTGAGGAAAC | Template DNA for *in vitro Cg-VPS51+DCTN1+SAC1* synthesis |
| *CgVPS51*-dsRNA-T7-R | GCGCTTGTGAAGCTGGTCGCGCAGCTCCTCTGACGCGGAG | |
| *CgDCTN1*-dsRNA-T7-F | CTCCGCGTCAGAGGAGCTGCGCGACCAGCTTCACAAGCGC | |
| *CgDCTN1*-dsRNA-T7-R | GTCCAACTTGTCGCGGAGATCGGCGTAAGACTCGCTCAC | |
| *CgSAC1*-dsRNA-T7-F | GTGAGCGAGTCTTACGCCGATCTCCGCGACAAGTTGGAC | |
| *CgSAC1*-dsRNA-T7-R | TAATACGACTCACTATAGGGAGAGTCAACCTGAGCAGCAAGATC | |
| *Cg-Actin*-qRT-F | CTCAGAGGCCTGCTGAAGATC | qRT-PCR Gene Expression |
| *Cg-Actin*-qRT-R | GCTTGGATTGATGTGTACAGTGC | |
| *CgDCL1*-qRT-F | TACTGACACGACGATTT | qRT-PCR *CgDCL1* Expression |
| *CgDCL1*-qRT-R | TTGAAGTTCCTCCCAC | |
| *CgDCL2*-qRT-F | CCGAACCTCAAATCC | qRT-PCR *CgDCL2* Expression |
| *CgDCL2*-qRT-R | CCGCCTCTTCAATCT | |
| *CgVPS51*-qRT-F | GACACTCACGTCCTCCG | qRT-PCR *CgVPS51* Expression |
| *CgVPS51*-qRT-R | TTGTCCTTCCGCATCT | |
| *CgDCTN1*-qRT-F | GAAGGCTATCATCCACTC | qRT-PCR *CgDCTN1* Expression |
| *CgDCTN1*-qRT-R | CTACGCAGCATCTCCA | |
| *CgSAC1*-qRT-F | GAGCCGACCCTGTTA | qRT-PCR *CgSAC1* Expression |
| *CgSAC1*-qRT-R | ACTAGAGGCCGCATT | |
| *RsPG*-dsRNA-T7-F | TAATACGACTCACTATAGGGAGATGACAGCAAGTACAGGT | Template DNA for *in vitro Rs-PG* synthesis |
| *RsPG*-dsRNA-T7-R | TAATACGACTCACTATAGGGAGATCACACTGGTGCCCGAA | |
| *RsDCTN1*-dsRNA-T7-F | TAATACGACTCACTATAGGGAGAGTTGCCTCGCACTAC | Template DNA for *in vitro Rs-DCTN1+SAC1* synthesis |
| *RsDCTN1*-dsRNA-T7-R | CCCACTTCAAGTCGGTCGCAGTCACACCAA | |
| *RsSAC1*-dsRNA-T7-F | TTGGTGTGACTGCGACCGACTTGAAGTGGG | |
| *RsSAC1*-dsRNA-T7-R | TAATACGACTCACTATAGGGAGACCGCCTGAGTCTGTA | |
| *Rs-Actin*-qRT-F | CCAACCGAGAAAAGATGACGC | qRT-PCR Gene Expression |
| *Rs-Actin*-qRT-R | CGTAAATTGGAACCGTATGCG | |
| *Os18S-rRNA*-qRT-F | ATGATAACTCGACGGATCGC | qRT-PCR Gene Expression |
| *Os18S-rRNA*-qRT-R | CTTGGATGTGGTAGCCGTTT | |
| *RsPG*-qRT-F | CGCCGAAGTGGATGT | qRT-PCR *RsPG* Expression |
| *RsPG*-qRT-R | TTGGACGAATTTCAGATAG | |
| *RsDCTN1*-qRT-F | TAGGGAGGGTGCTGT | qRT-PCR *RsDCTN1* Expression |
| *RsDCTN1*-qRT-R | AAATCTGTCTCCTGCTTC | |
| *RsSAC1*-qRT-F | CAATGATGCCACTCC | qRT-PCR *RsSAC1* Expression |
| *RsSAC1*-qRT-R | ATCTCGGCTCGTTAG | |

| | | |
|---|---|---|
| *AnpgxB*-dsRNA-T7-F | TAATACGACTCACTATAGGGAGACGACGATGTGTCCAGTGACTTC | Template DNA for *in vitro* *AnpgxB* synthesis |
| *AnpgxB*-dsRNA-T7-R | TAATACGACTCACTATAGGGAGAGCAGGACCACCGAAATCAGTATC | |
| *AnVPS51*-dsRNA-T7-F | TAATACGACTCACTATAGGGAGAGCCATGATGAAGATGCCGC | Template DNA for *in vitro* *An-VPS51+DCTN1+SAC1* synthesis |
| *AnVPS51*-dsRNA-T7-R | GACCGGACGAGCGCTTGGTGACGGACTTCGCCGAAGGGACGC | |
| *AnDCTN1*-dsRNA-T7-F | GCGTCCCTTCGGCGAAGTCCGTCACCAAGCGCTCGTCCGGTC | |
| *AnDCTN1*-dsRNA-T7-R | CTCCAGATGCTTCACTAGGCGGTCAAGGTGAGCTTGGACGGC | |
| *AnSAC1*-dsRNA-T7-F | GCCGTCCAAGCTCACCTTGACCGCCTAGTGAAGCATCTGGAG | |
| *AnSAC1*-dsRNA-T7-R | TAATACGACTCACTATAGGGAGACGATGTTGATGCCCTCGC | |
| *An-Actin*-qRT-F | ACGCTTGGACTGTGCCTC | qRT-PCR Gene Expression |
| *An-Actin*-qRT-R | CAATGGTTCGGGTATGTGC | |
| *AnpgxB*-qRT-F | TTCGGCGAGAAGCAGTCA | qRT-PCR *AnpgxB* Expression |
| *AnpgxB*-qRT-R | CACCGATAATGCAACCCG | |
| *AnVPS51*-qRT-F | CACAAACTGGCTGGAC | qRT-PCR *AnVPS51* Expression |
| *AnVPS51*-qRT-R | GCCTTGACCGAATACA | |
| *AnDCTN1*-qRT-F | GACGAGCAAGCACATAC | qRT-PCR *AnDCTN1* Expression |
| *AnDCTN1*-qRT-R | GCAGCGATCTCCAAC | |
| *AnSAC1*-qRT-F | CCCATTTATGCCGTGAC | qRT-PCR *AnSAC1* Expression |
| *AnSAC1*-qRT-R | GTGCCAACTTCAGTGTCG | |
| *VdDCL1*-dsRNA-T7-F | TAATACGACTCACTATAGGGAGATCATGACCACAGTCATGAG | Template DNA for *in vitro* *Vd-DCL1/2* synthesis |
| *VdDCL1*-dsRNA-T7-R | GTCGCCTGTGGCAGTAGATCGTTGACCTTCTGTGTATGG | |
| *VdDCL2*-dsRNA-T7-F | CCATACACAGAAGGTCAACGATCTACTGCCACAGGCGAC | |
| *VdDCL2*-dsRNA-T7-R | TAATACGACTCACTATAGGGAGACCAGTCGAACCTCGACAATG | |
| *VdDCTN1*-dsRNA-T7-F | TAATACGACTCACTATAGGGAGACTCGTCACGCCTGGCGCC | Template DNA for *in vitro* *Vd-DCTN1+SAC1* synthesis |
| *VdDCTN1*-dsRNA-T7-R | CTTGGTCCTCATGGCATCCAGCAGGCTGGCGGCGCGAGGCGTC | |
| *VdSAC1*-dsRNA-T7-F | GACGCCTCGCGCCGCCAGCCTGCTGGATGCCATGAGGACCAAG | |
| *VdSAC1*-dsRNA-T7-R | TAATACGACTCACTATAGGGAGAGCGGATTGCTGGTCTACTTGCATG | |
| *At-actin2*- qRT-F | ACCTTGCTGGACGTGACCTTACTGAT | qRT-PCR Gene Expression |
| *At-actin2*- qRT-R | GTTGTCTCGTGGATTCCAGCAGCTT | |
| *Vd-actin*-qRT-F | TAACTCTCCCGCTATGTATGTCGC | qRT-PCR Gene Expression |
| *Vd-actin*-qRT-R | GAGAGAAACCCTCGTAGATTGGC | |
| *VdDCL1*-qRT-F | GTCGCAGACCTTCCATACTT | qRT-PCR *VdDCL1* Expression |
| *VdDCL1*-qRT-R | AGATCTGTCCGAATACCTCTGAG | |

| | | |
|---|---|---|
| *VdDCL2*-qRT-F | GACTCTCCAGGCTTTCAAGA | qRT-PCR *VdDCL2* Expression |
| *VdDCL2*-qRT-R | TCTGAGAAATTGAGGTCACGAA | |
| *VdDCTN1*-qRT-F | CGTTGGCTCTGTTTCCT | qRT-PCR *VdDCTN1* Expression |
| *VdDCTN1*-qRT-R | GGTTCTCCATCAGCGTCT | |
| *VdSAC1*-qRT-F | TTCTGGAACCGAAATAC | qRT-PCR *VdSAC1* Expression |
| *VdSAC1*-qRT-R | CGGAGATGTTGGAGC | |
| *PiDCL1*-dsRNA -T7-F | TAATACGACTCACTATAGGGAGA GAAGTTCGTGGAGCTAGAGC | Template DNA for in vitro *PiDCL1* synthesis |
| *PiDCL1*-dsRNA -T7-R | TAATACGACTCACTATAGGGAGA GCCTTCTTTACCAGCGCTATC | |
| *PiHMP1*-dsRNA -T7-F | TAATACGACTCACTATAGGGAGA GGCTCCGGCCGACATCCCG | Template DNA for in vitro *PiHMP1* synthesis |
| *PiHMP1*-dsRNA -T7-R | TAATACGACTCACTATAGGGAGA GTGTCTTTAGTAAAGTCACCG | |
| *PiGPB1*-dsRNA -T7-F | TAATACGACTCACTATAGGGAGA ATGGGCGACGCTGCGGAGC | Template DNA for in vitro *PiGPB1* synthesis |
| *PiGPB1*-dsRNA -T7-R | TAATACGACTCACTATAGGGAGA CGACATTTGGGTGGGGCCAG | |
| *PiDCTN1*-dsRNA-T7-F | TAATACGACTCACTATAGGGAGA CAGCTTCAGCAGCAACAACAG | Template DNA for *in vitro Pi-DCTN1+SAC1* synthesis |
| *PiDCTN1*-dsRNA-T7-R | GTCACGACAGTATCAGAACGCG AAGAAGTAGGTTCCGCTGC | |
| *PiSAC1*-dsRNA-T7-F | GCAGCGGAACCTACTTCTTCGCG TTCTGATACTGTCGTGAC | |
| *PiSAC1*-dsRNA-T7-R | TAATACGACTCACTATAGGGAGA CGTGCACGTGCTGATTATCC | |
| *Pi-Tubulin*-qRT-F | GAATACGATCCCACGAA | qRT-PCR Gene Expression |
| *Pi-Tubulin*-qRT-R | AACGAGCCCAGTCCA | |
| *PiDCL1*-qRT-F | GCGTGTCGTGTTTGA | qRT-PCR *PiDCL1* Expression |
| *PiDCL1*-qRT-R | TTTCAGCACGTCAATT | |
| *PiGPB1*-qRT-F | ATTTCTTCCCGAGTGG | qRT-PCR *PiGPB1* Expression |
| *PiGPB1*-qRT-R | TTGTAGTCGTCGTAGCC | |
| *PiHMP1*-qRT-F | GTTTCCGCCGTCGTT | qRT-PCR *PiHMP1* Expression |
| *PiHMP1*-qRT-R | TCCGTGTCCGTAGTCG | |
| *PiDCTN1*-qRT-F | CTCTGTTTCCCAAGCG | qRT-PCR *PiDCTN1* Expression |
| *PiDCTN1*-qRT-R | ATCGCCCACATCCAA | |
| *PiSAC1*-qRT-F | CAGTAGTAGTGGGAGCG | qRT-PCR *PiSAC1* Expression |
| *PiSAC1*-qRT-R | GAGCGACACTTTGAGC | |

**Table S2.** The sequences of dsRNAs in this study.

| dsRNA | dsRNA sequence |
|---|---|
| *YFP*-dsRNA | TAATACGACTCACTATAGGGAGAATGGTGAGCAAGGGCGAGGAGCTGT<br>TCACCGGGGTGGTGCCCATCCTGGTCGAGCTGGACGGCGACGTAAACGG<br>CCACAAGTTCAGCGTGTCCGGCGAGGGCGAGGGCGATGCCACCTACGG<br>CAAGCTGACCCTGAAGTTCATCTGCACCACCGGCAAGCTGCCCGTGCCC<br>TGGCCCACCCTCGTGACCACCTTCGGCTACGGCGTGCAGTGCTTCGCCC<br>GCTACCCCGACCACATGCGCCAGCACGACTTCTTCAAGTCCGCCATGCC<br>CGAAGGCTACGTCCAGGAGCGCACCATCTTCTTCAAGGACGACGGCAAC<br>TACAAGACCCGCGCCGAGGTGAAGTTCGAGGGCGACACCCTGGTGAAC<br>CGCATCGAGCTGAAGGGCATCGACTTCAAGGAGGACGGCAACATCCTG<br>GGGCACAAGCTGGAGTACAACTACAACAGCCACAACGTCTATATCATG<br>GCCGACAAGCAGAAGAACGGCATCAAGGTGAACTTCAAGATCCGCCAC<br>AACATCGAGGACGGCAGCGTGCAGCTCGCCGACCACTACCAGCAGAAC<br>ACCCCCATCGGCGACGGCCCCGTGCTGCTGCCCGACAACCACTACCTGA<br>GCTACCAGTCCGCCCTGAGCAAAGACCCCAACGAGAAGCGCGATCACA<br>TGGTCCTGCTGGAGTTCGTGACCGCCGCCGGGATCACTCTCGGCATGGA<br>CGAGCTGTACAAGTAAAGAGGGATATCACTCAGCATAAT |
| *Ss-DCL1/2*-dsRNA | TAATACGACTCACTATAGGGAGACATTTCGGAGGCGGTCGAAACTCGCC<br>GGTTTCATGCGTGCCTCTCACTTCTTCAGTGAAGCTCGAAGGAACCACG<br>CTTGACCAAGTTAATATGTTCACCCTATGTCTATTTCAGGATGTGTTCAG<br>CAAAGCTTACAAATCGAATCCTGATAGTATGCCGTACTTTTTGGTTCCTA<br>TCAATTGTCCAAATCACACTGGGGACTGGAAAGATTATGAGCCCATGTC<br>AATAATTGATTGGGAAACTGTTCAATATGTCCAAGGCATGGCAAGAGGC<br>AGAACATAATCCTGGAATTGTTCGTACATCTATTATGAGCCTCGATGATT<br>TAAAGATTGAGCTCTGCCTTCCGATCGAGCCACCACCAATTCCGCCCTT<br>GACACTTCACTGGAATTCCAGCACAGAGCTGTCTGTCGAGTTTACAAAC<br>GATGCCGAAATAGGAACCACCGAGGACATGATATTACGAGAGGGATAT<br>CACTCAGCATAAT |
| *Ss-*<br>*VPS51+DCTN1+SAC1*<br>-dsRNA | TAATACGACTCACTATAGGGAGAGCATTGGCTGCGCTGAAGACGAAAC<br>CTTTATTGAAAGATGAAAGTATTCGAGATTTGGAGGGATTGAGATTAGA<br>TGTATGTGAGCGGTGGTTTGGTGATGAAATTCTTTACTTTACACCTTATG<br>TTCGACACGATGATTTGGAAGGATCATTAGCCGTTGAGACATTAAGAGG<br>TTGGGCGAAGAAAGCATCAGAGGTACTACTGGAAGGATTCACAAAGAC<br>TCTTCAAGGTCAATGCCAGAGATTGACGGTTACTCTTCAGCAGCGGAGG<br>ATTTCTGTGAATTGTGCGAGGCATCAGGTCATGATGTTCTACATTGCCCC<br>ATGTTCGGTCCTAATGGTAATAGTGGCAACTCTAGAGAGGAGTCTCCTA<br>AAGAGCAACGAACAGGAAAAGACGTTGTCATGGAAGGACTCAAACTAT<br>CACCCAAACGCTTACATTTCCAAGCAGGAAATCAATGGCCGCTTGACTG<br>AAATAATCATTAACAATACCGCTGTAAAATCTAGAGATAGAAAGTCCCA<br>TGTCTGTCAAAGCACCCTTGTAATCACGCTTTCTAGTACGAGTGAAGTCT<br>CCCTTCAGGGCTGCTGTAGAGGCATATTGCTTAGAGATAGCATCGCCAT<br>TATCAGAGAGGGATATCACTCAGCATAAT |
| *Bc-DCL1/2*-dsRNA<br>(Wang et al., 2016) | TAATACGACTCACTATAGGGAGATGCGGAAGAACTTGAAGGTTTGCTAC<br>ACAGTCAAATATGTACTGCAGAAGATCCCAGCTTGCTGCAGTACTCAAT<br>CAAAGGTAAACCTGAGACTCTTGCCTACTATGATCCCTTGGGCCCGAAA<br>TTCAATACTCCTCTTTATCTTCAAATGCTCCCGCTTCTAAAAGACAATCC<br>TATCTTTCGGAAGCCATTTGTATTTGGGACAGAAGCCAGTAGAACTCTA<br>GGATCTTGGTGTGTTGACCAGATCTGGACGGATGCCATTTGCTGCACGC<br>CAAAAATACATCGAGCAGATCTTCGCCTTCGAGTAAAGCTACCACTTCT<br>ATCTATTATCTACTATACCCCAGAGTCAAATATCATCGTGACGAAAACT<br>GTGGCGAGCCTGAGAAAGATTGTGCAAAGTCTCAACATTTTCGAAGACC<br>CCTACGTTTTGACACTAAAAAGGAGTGATAGCGAAAAAAGTCAACGTG<br>AGCTGGCGAAAGTACTCAAGAGTAGAGGGATATCACTCAGCATAAT |
| *Bc-*<br>*VPS51+DCTN1+SAC1*<br>-dsRNA | TAATACGACTCACTATAGGGAGATTCGTTCCAGGAGTTACACGCAAAGA<br>TCACTTCACTGGTGGACCAGCAAAAAGATAGTAAACATATCGGGAAAA<br>TATCGATATATATTCTCCGAATTCTACGAGATATCAGAGCAGAATTACC |

| | |
|---|---|
| | TAGTAACCCTGCACTACAAAAGTTTGGACTCTCACTTGTCTCATCATTCG<br>TGCTCTCGTCGAAAGCAAAGACAATAAAGAAGAACAAATGGAGCGCGA<br>GTTGGAGGGATTGCGAAGAGGAAGTGTTAGCAATCCTACTACGCATCGT<br>ACTAGTGCCATGAGCAGCGGAACTGTGACTCAGGATAGGAATTCTCTCC<br>AAGACAATAAGAGCACAGTTGTAAGCTGGACGTTGTTCAAAGTGCAGT<br>GGCAAAGCGAGCACTTGAAATGCAGTTAAAGAATGAGGGACTAGATGT<br>CACTCTACAAATTGATCAAACTCAACAATGGTTCAATACTTTGTGGGCC<br>GACAATGGTGACGCCATTTCTAAGCAATACGCTTCTACAGCAGCATTGA<br>AGGAGAGGGATATCACTCAGCATAAT |
| *Cg-DCL1/2*-dsRNA | TAATACGACTCACTATAGGGAGAGTCAGATGTCTATCCCGTTTCCTTGCC<br>CACTCCTATCCGGGTGTCTGCAGAAGACTCCGAAGCCCTGAAAGAGTTT<br>ACCTTACGCATTTTCAAAGACGTCTTCAGCAAGCAGTTCGAGGCCCCAA<br>TTACTGACGTGCCGTACTTTCTTGCCCCTGCCACATTCTATCACTCGGCA<br>AAGCTTTCTGAAGTCGGTGATCAGCTTCCCATCGAGGAGCCTGAGAACG<br>GAGACCATACTGCTGTACTTCTCGCCTTGCCTTACGGCCATCGCTGGTTG<br>TCGAGTCCGCTTCAGCAGGTAATCAGGTTTAGCGCAGTTGGTGCCGAAC<br>TTTCTCTGGATGGTATTGGAACGAAGCATTTTGTGCCAGGGGAGATGTC<br>TGCACACGACCTCACCTGCTTCAGAGGGATATCACTCAGCATAAT |
| *Vd-DCL1/2*-dsRNA (Wang et al., 2016) | TAATACGACTCACTATAGGGAGATCATGACCACAGTCATGAGTTCTCAC<br>CGAATGAAGACCCAGGGTCACTGATCGACTGGAGCCATCTGCTGTCGAC<br>CAAAGAGGTTGAGTACTTGCCTTGGGATGAAGATCACAGTCCCAGCTTC<br>TATCAAAGCAAGTTTGTGATTGATCCATACACAGAAGGTCAACGATCTA<br>CTGCCACAGGCGACAAACAAGAAGCCATATGCTAGGGTTTATCCGGCAC<br>CGTGGGCGAAAGTCGACACGATTCCATTAGATCATGCTTACTTTGGGGC<br>GTTGATCCCTTTCATTTCACACATTGTCGAGGTTCGACTGGAGAGGGAT<br>ATCACTCAGCATAAT |
| *PiHMP1*-dsRNA | TAATACGACTCACTATAGGGAGAGGCTCCGGCCGACATCCCGTCTTCAC<br>AAGACGAGAATCTCAAGAAACAGGAAGAGCGTGGCTTCTTTGACTGGTT<br>TGGCAATGGCGACGACACGCCTGCCCCGGCTGCTGACGACAACTCCGGT<br>AAATATGCGACTACGACACCGATCCCTGGCACGGCTGCGCGGCCGTCTA<br>CGAGTAGTCTGATGTCAAAGTATGGTAGTATGCTCGGTGACTTTACTAA<br>AGACACGAGAGGGATATCACTCAGCATAAT |
| *PiPGB1*-dsRNA | TAATACGACTCACTATAGGGAGAATGGGCGACGCTGCGGAGCTCAAGA<br>AGAAATGCGAGAGCCTCAAGGAGACGATTGAGAAGACGCGTGAAGCCA<br>AGAGTGATGGCGGCTTCCAGAGCGCCAATGCCAGCTCCGGCGCCAAGG<br>CGATCCTGGCCCCACCCAAATGTCGAGAGGGATATCACTCAGCATAAT |
| *PiDCL1*-dsRNA | TAATACGACTCACTATAGGGAGAGAAGTTCGTGGAGCTAGAGCTGCAG<br>GCCAATTTACAGGCGTGTATTGTGGACGACCCAGACAACGTGAACAACA<br>GGAAGAAGCGCAAGCAGCAAGAAGACGATCCTGATATCGACGATTCGA<br>TGATGGGAGAGGGCATTGATAGCGACTTCACGGAAGATGGAGCGGTGG<br>GTGAGGAAGAGGGCACGGACGAGATGATGGAGTGCAGCAATGCAGAA<br>GATAGCGCTGGTAAAGAAGGCAGAGGGATATCACTCAGCATAAT |

"_" representative T7 promoter sequence, color representative different genes sequence fragments that used for dsRNA templates.
